# Survival-associated cellular communication networks in human solid cancers

**DOI:** 10.64898/2026.10.16.759703

**Authors:** Derek Lee, Varchas Bharadwaj, Jingtong Liang, Georgia R. Lill, Jeong Hyun Ji, Emilie Campbell, Kenneth Ho, Timothy E. O’Sullivan

## Abstract

Single-cell atlases have enabled the prediction of human cell-cell communication (CCC) networks in cancer, but whether predicted communication networks have clinical significance is not well understood. Here, we integrated a pan-solid cancer scRNA-seq atlas comprising 78 cell types from 318,426 quality-controlled cells with bulk RNA-seq and clinical outcome data from The Cancer Genome Atlas (TCGA) to develop Single-cell-based Kaplan-Meier Analysis Network (SKyNet). SKyNet prioritizes ligand-receptor interactions associated with overall survival across 11 human solid cancers, predicting unique interactions not inferred by previous CCC algorithms. In pancreatic adenocarcinoma, SKyNet-predicted endothelial-pericyte interactions were associated with poor outcomes that were validated by independent spatial transcriptomics and survival cohorts. In lung adenocarcinoma, lymphatic endothelial cell interactions consisting of collagen-integrin pairs were associated with poorer survival outcomes. In kidney renal clear cell carcinoma and kidney renal papillary cell carcinoma, fibroblast communication networks displayed disparate survival associations, with the DSC2-DSG2 axis associated with better survival outcomes. Thus, SKyNet provides a hypothesis-generating framework for cell type-specific communication pathways associated with patient outcomes.

## INTRODUCTION

Single-cell RNA sequencing (scRNA-seq) has transformed our understanding of the human tumor microenvironment (TME) by resolving the diversity of cell types and states across cancers. Integrated pan-cancer tumor atlases have revealed both recurrent multicellular ecosystems and tissue-specific programs, further defining how cellular abundance and cell type-specific programs are rewired during tumorigenesis. Computational frameworks such as CellPhoneDB^1^, CellChat^2^, and NicheNet^3^ use ligand-receptor expression and downstream transcriptional programs to infer cell-cell communication (CCC) between defined cell populations. Together, these advances have moved the field beyond descriptive inventories of TME composition toward testable cellular communication circuits that may coordinate tumor progression or rejection.

However, communication networks predicted from cancer atlases could reflect homeostatic tissue communication networks or cellular responses to treatment, without revealing the importance of these pathways in tumor suppression or progression. Furthermore, clinical variables are often not incorporated into the prioritization of cellular communication^4^, and the relationships between inferred cellular communication networks and patient outcomes remain incompletely understood across human cancers. As metastatic disease is present in many patients at the time of death, and thromboembolism contributes to cancer mortality^5^, defining the cellular communication networks associated with survival in human solid tumors may reveal cell type-specific interactions relevant to patient outcomes that are not revealed from canonical CCC analyses.

To identify putative communication networks associated with clinical outcomes across human solid tumors, we developed a high-resolution solid cancer scRNA-seq atlas of healthy and cancerous tissue using patient-matched samples from seven common human solid cancers, comprising 78 unique cell types identified from 318,426 quality-controlled cells. We then developed <u>S</u>ingle-cell-based <u>K</u>aplan-Meier Anal<u>y</u>sis <u>Net</u>work (SKyNet), a computational framework that nominates clinically relevant cell communication interactions within the TME from publicly available bulk RNA-seq, single-cell RNA-seq, and clinical outcome data. We applied SKyNet to 11 different human cancers to identify CCC pathways associated with patient outcomes. SKyNet highlighted unique cancer-specific and tissue-independent CCC pathways associated with survival outcomes, emphasizing the importance of stromal cell interactions with extracellular matrix in driving poorer outcomes across a subset of human cancers. In case studies of pancreatic adenocarcinoma (PAAD), lung adenocarcinoma, and kidney renal cell cancers, SKyNet predicted CCC associated with poor and improved overall survival, many of which were supported by independent spatial transcriptomics datasets and/or previous research. SKyNet also identified new CCC pathways, such as the DSC2-DSG2 signaling axis, which was associated with prolonged survival in kidney renal clear cell carcinoma (KIRC). Overall, our analyses demonstrate SKyNet’s utility as a hypothesis-generation tool that can be broadly applied across cancer and other disease contexts to interrogate CCC pathways potentially implicated in disease progression and patient outcomes.

## RESULTS

### An integrated human solid tumor atlas identifies cell type-specific associations with survival across human cancers

To identify cell types present in human solid cancers at high resolution, we selected 6 existing large-scale scRNA-seq datasets representing 7 major human solid tumors from 64 patients: breast adenocarcinoma (BRCA), KIRC, head and neck squamous cell carcinoma (HNSC), colon adenocarcinoma (COAD), hepatocellular carcinoma (HCC), lung adenocarcinoma (LUAD), and lung squamous cell carcinoma (LUSC) (**Figure 1A**, **Table 1**). After quality control to exclude cells with low transcript abundance or high representation of mitochondrial transcripts, we obtained 318,426 cells. We then integrated these data across tumor types and patient samples using Harmony^6^, which allowed clustering by transcriptomic similarity rather than by patient or tumor type (**Figure S1A,B**). Unsupervised clustering identified three broad clusters (lymphoid, myeloid, and structural) that were sub-clustered to resolve cell types (**Figure 1B**). This multi-level clustering approach enabled high-resolution annotation of conserved cell types and discovery of tissue-specific and rare cell types, while limiting the inclusion of doublets and transitional cell types.

**Figure 1.**
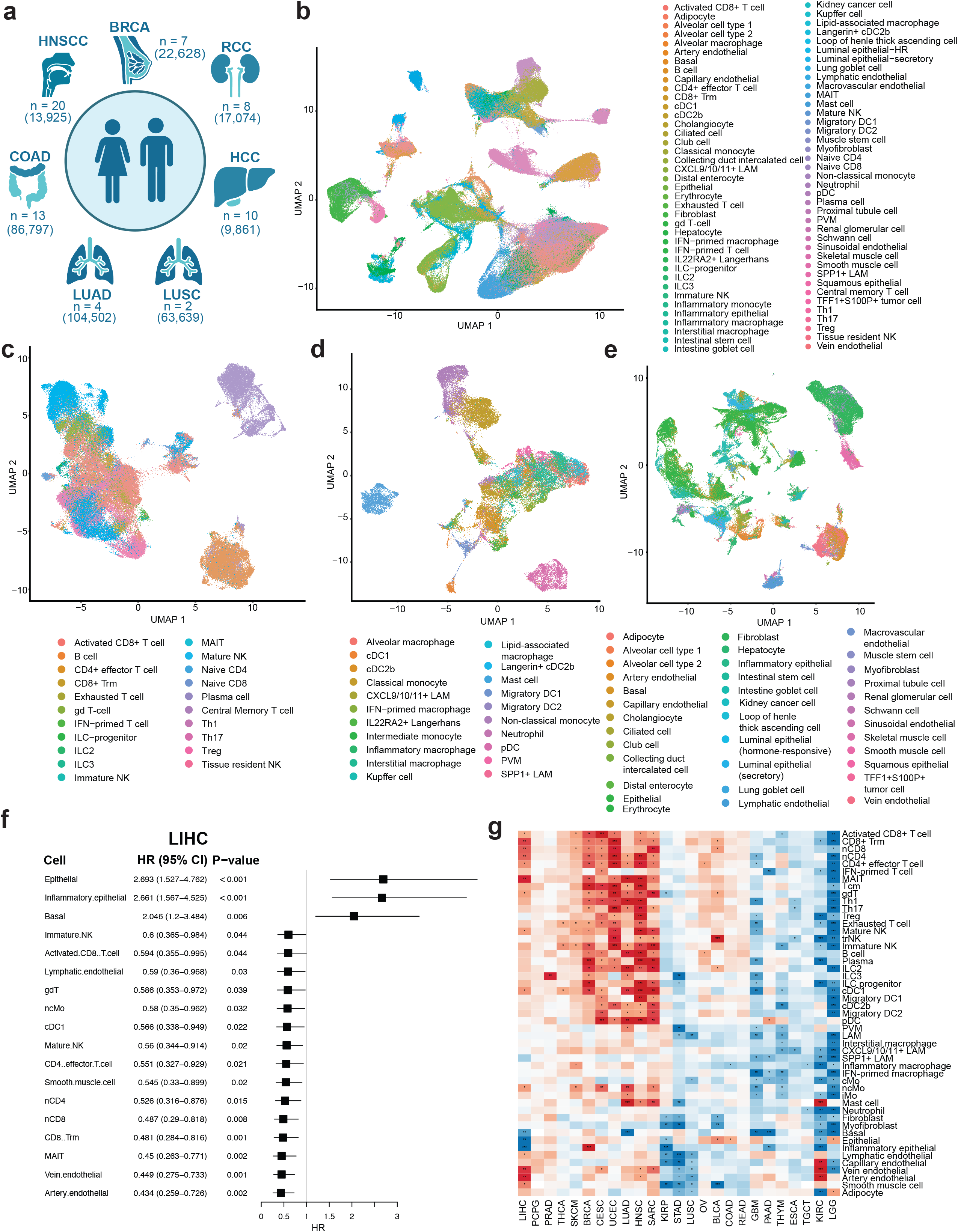
JadeCell: lineage markers delineating cell types from a solid cancer single-cell transcriptomics atlas. **(a)** Overview of patient population size and filtered cells in each scRNA-seq dataset of seven solid tumors of distinct origin in the integrated atlas (breast adenocarcinoma (BRCA), renal cell carcinoma (RCC), head and neck squamous cell carcinoma (HNSC), colon adenocarcinoma (COAD), hepatocellular carcinoma (HCC), lung adenocarcinoma (LUAD), and lung squamous cell carcinoma (LUSC)). **(b-e)** Uniform manifold approximation and projection (UMAP) plots of; all cell types **(b)**, lymphoid cells **(c)**, myeloid cells **(d)**, and stromal, parenchymal, and tissue-specific cells **(e)** across the integrated single-cell atlas. **(f)** Forest plot of associations between cell type abundance and overall survival (OS) among patients with liver hepatocellular carcinoma (LIHC) from The Cancer Genome Atlas (TCGA) using Cox proportional hazard models; positive hazard ratios indicate greater risk of event (death) associated with abundance of given cell type. **(g)** Heatmap of associations between cell type abundance and OS across cell types in 25 solid tumors from TCGA; red indicates cell types whose abundance was associated with longer OS, while blue represents those whose abundance was associated with shorter OS; tumors are clustered via hierarchical clustering of survival differences across cell types. *p < 0.05, **p < 0.01, ***p < 0.001; ****p < 0.0001; ns, not significant.

**Table 1.**
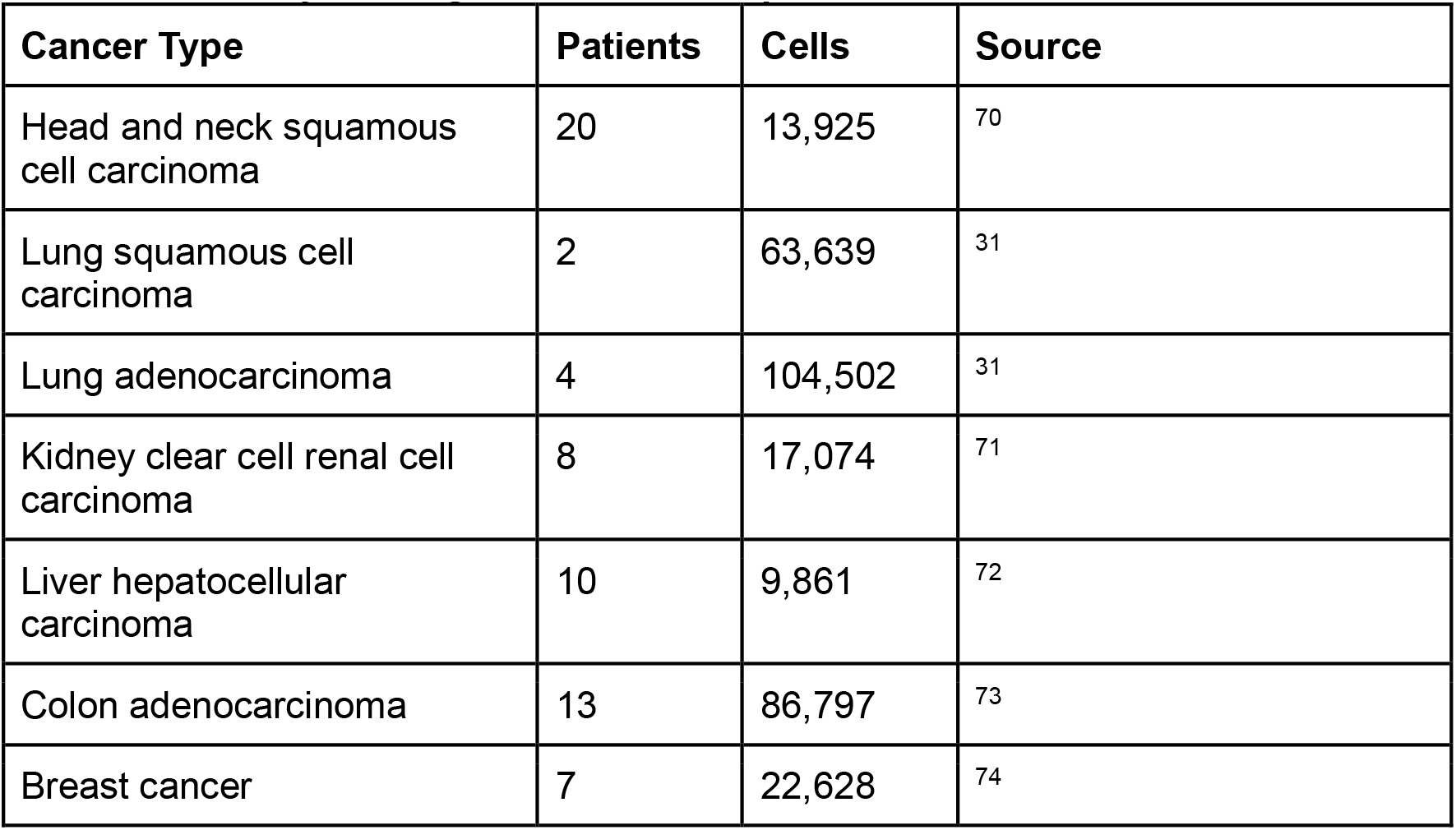
Summary of integrated scRNA-seq datasets.

| **Cancer Type** | **Patients** | **Cells** | **Source** |
| --- | --- | --- | --- |
| Head and neck squamous cell carcinoma | 20 | 13,925 | <sup>70</sup> |
| Lung squamous cell carcinoma | 2 | 63,639 | <sup>31</sup> |
| Lung adenocarcinoma | 4 | 104,502 | <sup>31</sup> |
| Kidney clear cell renal cell carcinoma | 8 | 17,074 | <sup>71</sup> |
| Liver hepatocellular carcinoma | 10 | 9,861 | <sup>72</sup> |
| Colon adenocarcinoma | 13 | 86,797 | <sup>73</sup> |
| Breast cancer | 7 | 22,628 | <sup>74</sup> |

Our integrated cell atlas resolved 21 cell types from the lymphoid lineage, including T cells, natural killer (NK) cells, B cells, plasma cells, and innate lymphocytes (**Figure 1C**). Within the T cell cluster, we identified naïve, helper, cytotoxic, memory, exhausted, and unconventional T cell subsets with high granularity. For instance, we detected a population of mucosal-associated invariant T (MAIT) cells characterized by *KLRB1* and *SLC4A10* expression^7^. Myeloid cells were clustered into 21 subtypes of macrophages, monocytes, dendritic cells (DCs), mast cells, and neutrophils (**Figure 1D**). The macrophage population consisted of tissue-resident cell types, including Kupffer cells and alveolar macrophages; tumor-associated macrophages with elevated expression of *SPP1* and angiogenesis-related genes such as *VEGFA, THBS1*, and *VCAN*; and a population of *TREM2^+^*lipid-associated macrophages (LAM) enriched for *CXCL9, CXCL10, and CXCL11,* which have been shown to play an important role in stimulating the migration and infiltration of immune cells into the solid TME^8,9^. Within the dendritic cell population, we identified a group of type 2 classical dendritic cells (cDC2b) enriched for *CD207* (langerin) and *CD1A*, consistent with mucosal Langerhans cells^10^. Additionally, we identified a subpopulation of Langerhans cells with elevated expression of *IL22RA2*. Structural cells were sub-clustered within each tissue independently, identifying 36 cell types including tissue-specific cells (such as lung and intestinal goblet cells, alveolar cells, and renal glomerular cells), fibroblasts, endothelial cells, epithelial cells, and smooth muscle cells, as well as rare cell types such as adipocytes and erythrocytes (**Figure 1E**). In LUAD, we identified a subpopulation of epithelial cells enriched for *TFF1, S100P, and TFF3*, which have been previously associated with poor outcomes and immunotherapy response in lung cancer^11^. In total, our integrated cell atlas identified 78 cell types across 7 distinct tumor types (**Supplemental Table 1**).

Given the high resolution of our integrated cell atlas, we created a set of cell type-defining gene modules called JadeCell by performing differential gene expression analysis on our atlas for each cell type and further curating a 20-gene list of differentially expressed genes in addition to canonical genes from existing atlases (**Supplemental Table 2**). When applied to the integrated atlas, JadeCell modules distinguished cell types based on Seurat module scores with high precision (**Figure S1C**). JadeCell modules also achieved higher mean area under the precision-recall curve than the gene modules from xCell^12^ and CellTypist^13^ across human solid tumor scRNA-seq datasets, both within and independent of our cell atlas (**Figure S1D, Table 2**). To better understand the role of these identified cell types in human solid cancers, we used JadeCell to identify relationships between cell abundance and patient outcomes. JadeCell gene modules were used to deconvolute cell type abundance in bulk RNA-seq data from the Cancer Genome Atlas (TCGA)^14^, and hazard ratios for each cell type were calculated using a univariate Cox proportional hazards model. In the TCGA liver hepatocellular carcinoma (LIHC) cohort, enrichment of epithelial, inflammatory epithelial, and basal cells was associated with significantly shorter overall survival, whereas several immune and endothelial cell types were associated with prolonged survival (**Figure 1F**). These findings align with previous research showing that high endothelial venules are associated with tumor-infiltrating lymphocyte recruitment^15^. Similarly, a subset of LIHC cases are driven by cytokeratin (CK) 19-positive cancer cells expressing a strong basal cell-like phenotype correlated with worse survival outcomes^16^. To evaluate whether the association between cell types and survival varied across tumor types, we applied this analysis to 25 solid tumors from TCGA using cell types shared across tissues identified in our cell atlas, revealing three broad categories of cell type abundance associations with overall survival (**Figure 1G**). One cluster, including LIHC, head and neck squamous cell carcinoma (HNSC), skin cutaneous melanoma (SKCM), cervical squamous cell carcinoma and endocervical adenocarcinoma (CESC), and uterine corpus endometrial carcinoma (UCEC), showed a favorable prognosis with higher abundance of lymphocytes, consistent with the reported responsiveness of these cancers to immune checkpoint blockade^17^. A second cluster, including kidney renal clear cell carcinoma (KIRC), lung squamous cell carcinoma (LUSC), and colon adenocarcinoma (COAD), showed no strong association between immune cells and outcomes, but endothelial cells were associated with poorer survival. The third cluster, including glioblastoma (GBM), pancreatic adenocarcinoma (PAAD), and brain lower grade glioma (LGG), showed negative associations across all immune cells but a favorable prognosis with endothelial cells, consistent with the poor response of these cancers to checkpoint blockade^18,19^. Thus, JadeCell gene modules can discriminate cell type associations with patient outcome across cancers in agreement with previous findings.

**Table 2.**
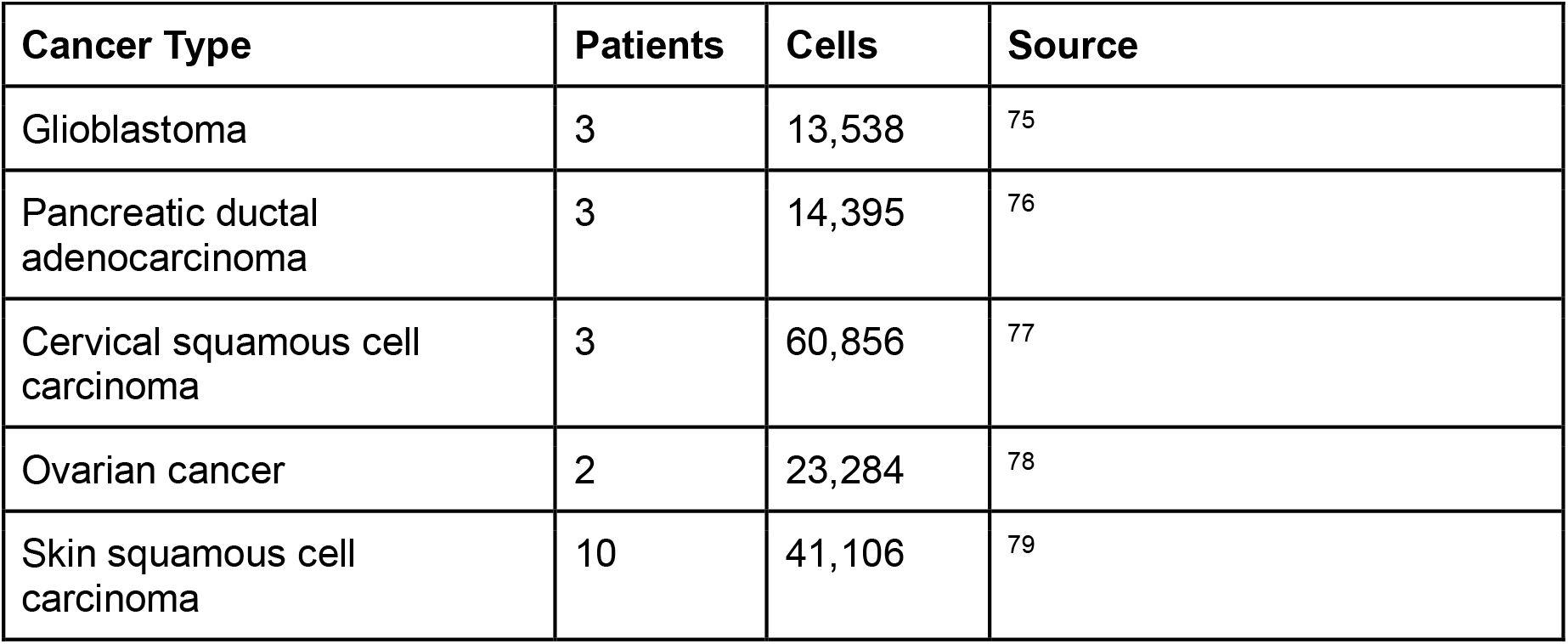
Summary of individual scRNA-seq datasets.

| **Cancer Type** | **Patients** | **Cells** | **Source** |
| --- | --- | --- | --- |
| Glioblastoma | 3 | 13,538 | <sup>75</sup> |
| Pancreatic ductal adenocarcinoma | 3 | 14,395 | <sup>76</sup> |
| Cervical squamous cell carcinoma | 3 | 60,856 | <sup>77</sup> |
| Ovarian cancer | 2 | 23,284 | <sup>78</sup> |
| Skin squamous cell carcinoma | 10 | 41,106 | <sup>79</sup> |

### Single-cell-based Kaplan-Meier Analysis Network (SKyNet)

To facilitate the discovery of clinically relevant CCC within the TME, we developed SKyNet, a cell-communication prediction algorithm prioritized by patient outcomes. First, SKyNet integrates patient-matched bulk RNA-seq and clinical outcome data from TCGA (**Figure S2, Supplemental Table 3**). SKyNet then stratifies patients based on top and bottom quartile expression of predefined JadeCell lineage markers for a given cell type, termed “High” and “Low”, and uses Kaplan-Meier analysis to assess differences in overall survival between the two strata (**Figure 2A**). To evaluate survival differences associated with ligand or receptor expression, patients with cell populations defined as “High” by cell-specific lineage markers were then further stratified based on ligand or receptor expression, by top and bottom quartiles. SKyNet then uses Kaplan-Meier analysis to compare overall survival between the "Cell High, Ligand/Receptor High" and "Cell High, Ligand/Receptor Low" strata (**Figure 2B**). To extend this framework to CCC, SKyNet queries the scDiffCom database of human ligand-receptor pairs^20^ and uses solid tumor scRNA-seq data of the same tumor type to identify the cell types that express each ligand and receptor (**Figure 2C**). SKyNet then applies Kaplan-Meier analysis to all possible pairs of ligand-expressing cells (senders) and ligand-receiving cells (receivers) (**Figure 2D**). Specifically, SKyNet computes the survival differences associated with ligand or receptor expression by top and bottom quartiles within each cell type. Using these survival differences together with scRNA-seq data that designate cell types as senders and/or receivers, SKyNet identifies sender-ligand and receiver-receptor pairs that are jointly associated with significant differences in overall survival. Finally, SKyNet removes nonspecific biomarkers and interactions with incongruent survival associations between the three tested combinations (sender and ligand, receiver and ligand, and receiver and receptor) before using permutation testing to identify statistically robust combinations (**Figure 2E**).

**Figure 2.**
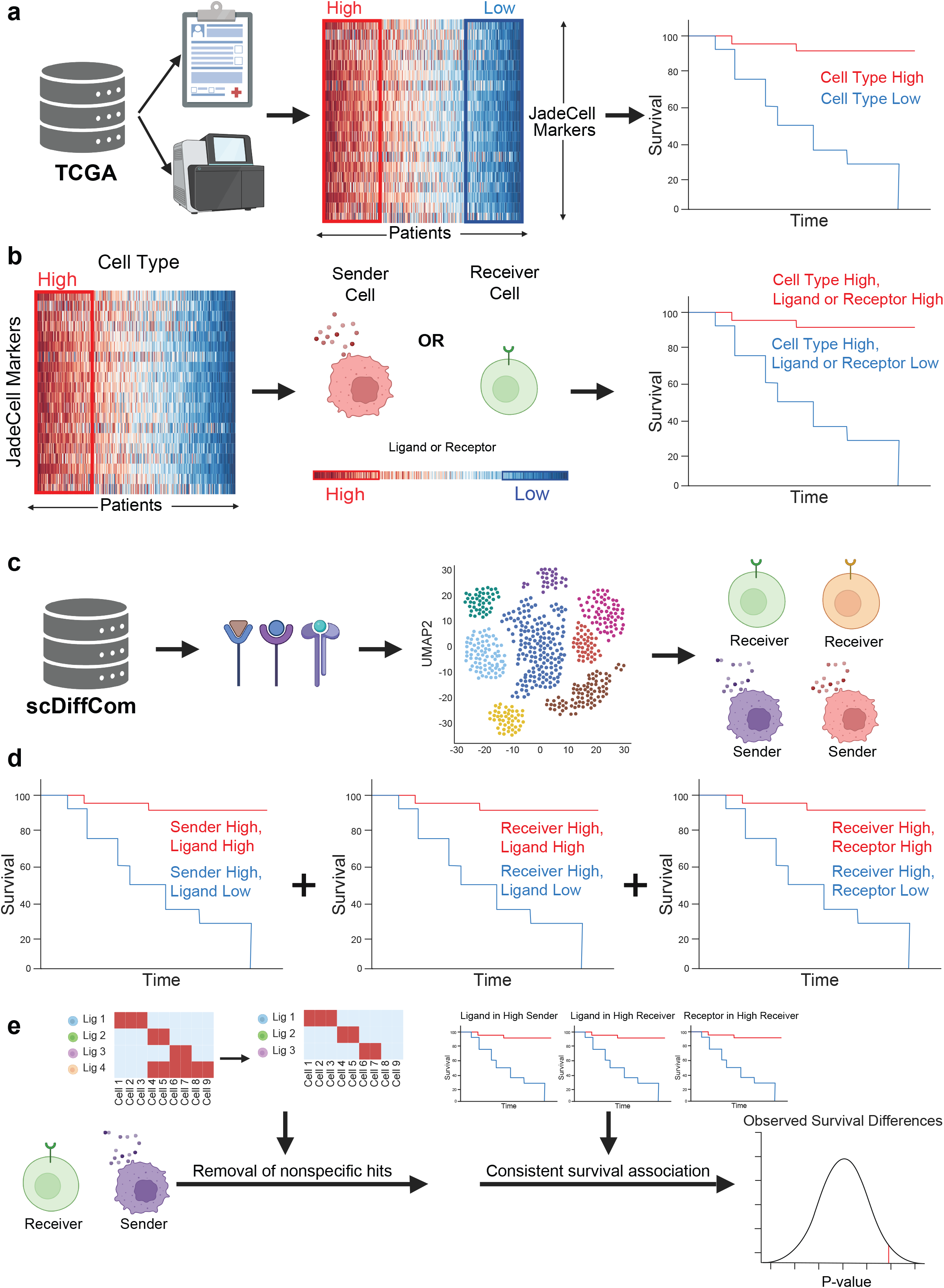
Schematic of SKyNet: a framework to infer clinically relevant intercellular ligand-receptor communications. **(a)** Schematic of patient outcomes based on cell type enrichment across solid cancers from The Cancer Genome Atlas (TCGA). **(b)** Schematic of patient outcomes based on cell type and ligand/receptor enrichment across solid cancers from TCGA. **(c)** Schematic of Human ligand-receptor pairs based on the scDiffCom database were used to identify cell type-specific ligand and receptor expression based on publicly available scRNA-seq datasets. **(d)** Schematic of survival analysis among TCGA patients for all possible sender-ligand, receiver-ligand, and receiver-receptor pairs within a given cancer. **(e)** Removal of nonspecific hits defined as significant across at least two-thirds of cell types, confirmation of concordance between sender-ligand, receiver-ligand, and receiver-receptor survival association, and prioritization of sender-ligand and receiver-receptor pairs based on permutation testing of survival trends. Figure created with BioRender.

### SKyNet identifies ligand-receptor interactions associated with overall survival across human solid cancers

To determine whether SKyNet CCC results were distinct from widely used CCC algorithms, we applied SKyNet to 11 different human solid cancers from TCGA and compared its outputs to CellChat^2^ and NicheNet^3^ (**Supplemental Table 4; Figure 3A**). Across 11 high-quality human solid cancer scRNA-seq atlases, CellChat predicted the most interactions while SKyNet predicted the fewest, although this trend was not always consistent within each cancer (**Figure 3B**). Moreover, the three algorithms each predicted a different distribution of CCC partners when each sender and receiver was categorized as immune or stromal (**Figure 3C**). While immune-immune interactions made up the largest fraction of predicted CCC in all three networks, CellChat predicted a greater proportion of stromal-immune interactions, while NicheNet predicted a greater proportion of immune-stromal interactions and SKyNet stromal-stromal interactions. Only one ligand-receptor interaction was predicted by all three algorithms, with 4 predicted by both SKyNet and NicheNet and 8 predicted by both CellChat and SKyNet (**Figure 3D**). Interestingly, this was not due solely to differences in the ligand-receptor databases used by each algorithm; the SKyNet database shared 52.8% of its interactions with the NicheNet database, 35.0% of its interactions with the CellChat database, and 14.7% of its interactions with both (**Figure S3A**). Finally, we used Biomni^21^ to categorize predicted ligand-receptor interactions by their function (**Figure 3E**, **Figure S3B,C, Supplemental Table 5**). Hierarchical clustering within SKyNet-predicted interactions revealed patterns in interaction families and survival associations across cancers. LUAD, LUSC, PAAD, and KIRP patients demonstrated similar negative survival associations with collagen, laminin, and extracellular matrix glycoprotein signaling. In general, extracellular matrix glycoprotein interactions were broadly associated with poorer survival outcomes but were associated with positive survival in KIRC. Meanwhile, chemokines and cell adhesion interactions in KIRC and CESC patients were associated with positive survival outcomes but showed absent or negative associations in other cancer types. The rest of the cancers analyzed (OV, COAD, HNSC, LIHC, and GBM) had fewer interactions associated with survival outcomes. When comparing communication networks, CellChat interactions concentrated on a narrower subset of families, with most interactions related to MHC, collagen, and ECM glycoproteins. NicheNet predicted similar proportions of each interaction family across cancers, with hierarchical clustering unable to distinguish observable clusters. NicheNet also had greater representation of TGF-beta and TNF superfamily interactions than CellChat or SKyNet. Together, these data suggest that SKyNet predicts a distinct CCC network from traditionally used algorithms.

**Figure 3.**
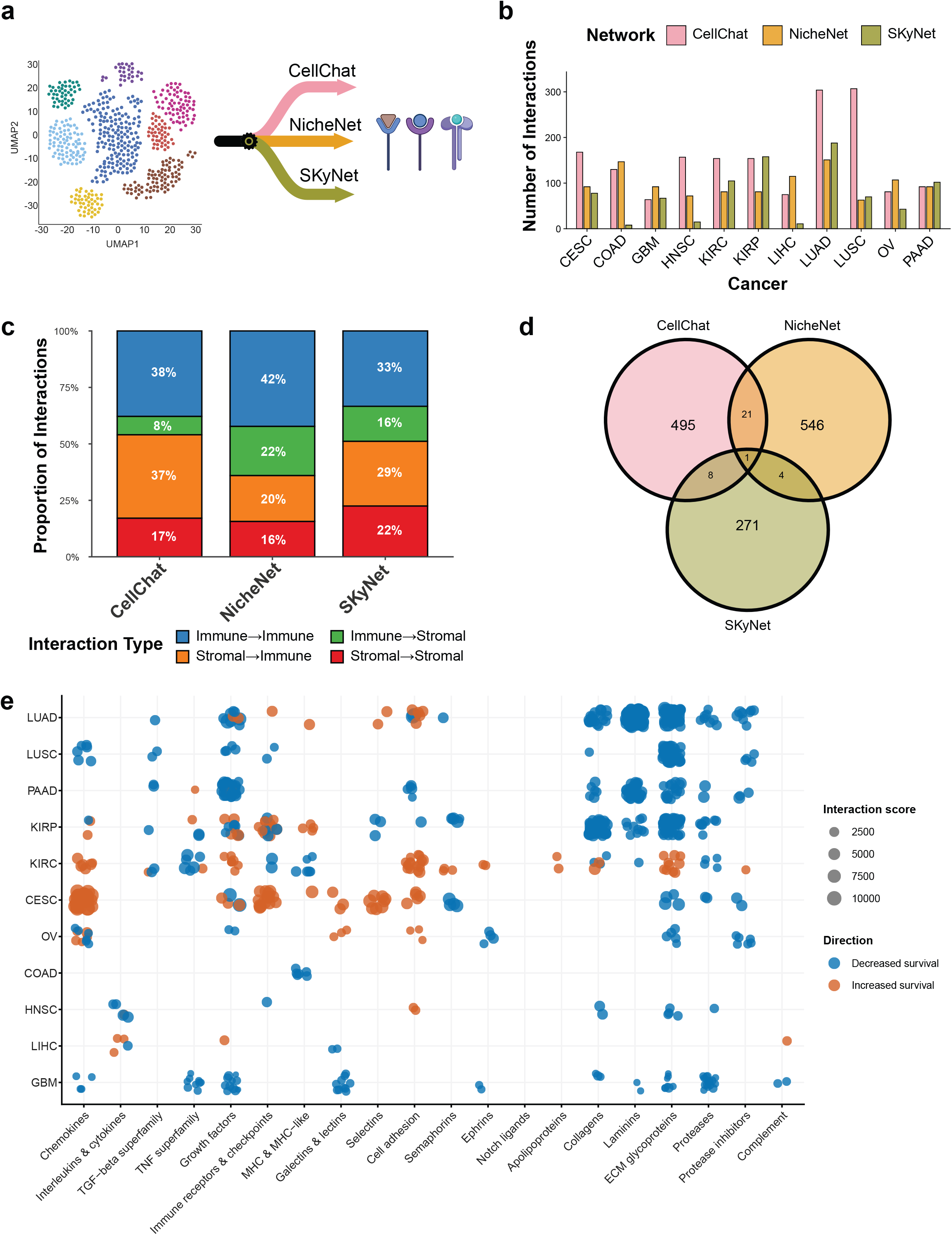
Survival-based communication network analysis identifies distinct ligand-receptor interactions across cancer. **(a)** Schematic of CCC inference performed via CellChat, NicheNet and SKyNET algorithms. **(b)** Number of significant ligand–receptor interactions identified by each algorithm in each tumor type. **(c)** Proportion of interactions assigned to each sender-receiver class (immune-immune, immune-stromal, stromal-immune, stromal-stromal), pooled across all tumor types. **(d)** Venn diagram of unique ligand–receptor pairs identified by each algorithm, pooled across all tumor types. **(e)** Functional classification of SKyNet interactions across cancers. Each ligand–receptor pair was assigned to one of 16 functional classes; point size denotes interaction score, and color denotes the direction of the survival association (blue, decreased survival; orange, increased survival).

### SKyNet-predicted endothelial interactions in pancreatic adenocarcinoma are supported by independent spatial and clinical cohorts

To assess the ability of SKyNet to identify cell types and pathways associated with overall survival, we applied it to the TCGA pancreatic adenocarcinoma (PAAD) dataset. SKyNet predicted 102 ligand-receptor interactions significantly associated with differences in patient outcomes, almost all of which were negatively associated with survival (**Figure S4A, Supplemental Table 4**). Endothelial cell interactions were associated with poor overall survival: vein endothelial cells were frequently predicted as a hub of outgoing ligands, whereas artery and capillary endothelial cells were frequently predicted to be receiving cell populations (**Figure 4A,B**). To verify whether these predicted interactions occur in patients, we utilized a publicly available Xenium spatial transcriptomics dataset of six PAAD patients^22^. Of the unique ligand-receptor pairs associated with SKyNet-predicted endothelial interactions, 25 of 40 (62.5%) showed co-localization by Moran’s I analysis^23^ (**Figure S4B**). For example, for the interaction between WNT5A from vein endothelial cells and LDLR from pericytes, both the ligand-receptor and sender-receiver combinations exhibited spatial colocalization (**Figure 4C**). This corroborates previous research indicating Wnt activation as a potential driver of PAAD proliferation and differentiation^24^. Additionally, several of the SKyNet-predicted endothelial interactions involved laminin-integrin interactions, which promote key steps of cancer metastasis like adhesion and migration and may therefore be linked with patient outcomes^25^. Consistent with this possibility, LAMA3 expression has been associated with increased liver metastasis, perineural invasion, and poor overall survival in pancreatic adenocarcinoma^26,27^. Of these laminin interactions, LAMA3 from pericytes to ITGA6/ITGB1 on artery endothelial cells had the highest SKyNet-predicted interaction score.

**Figure 4.**
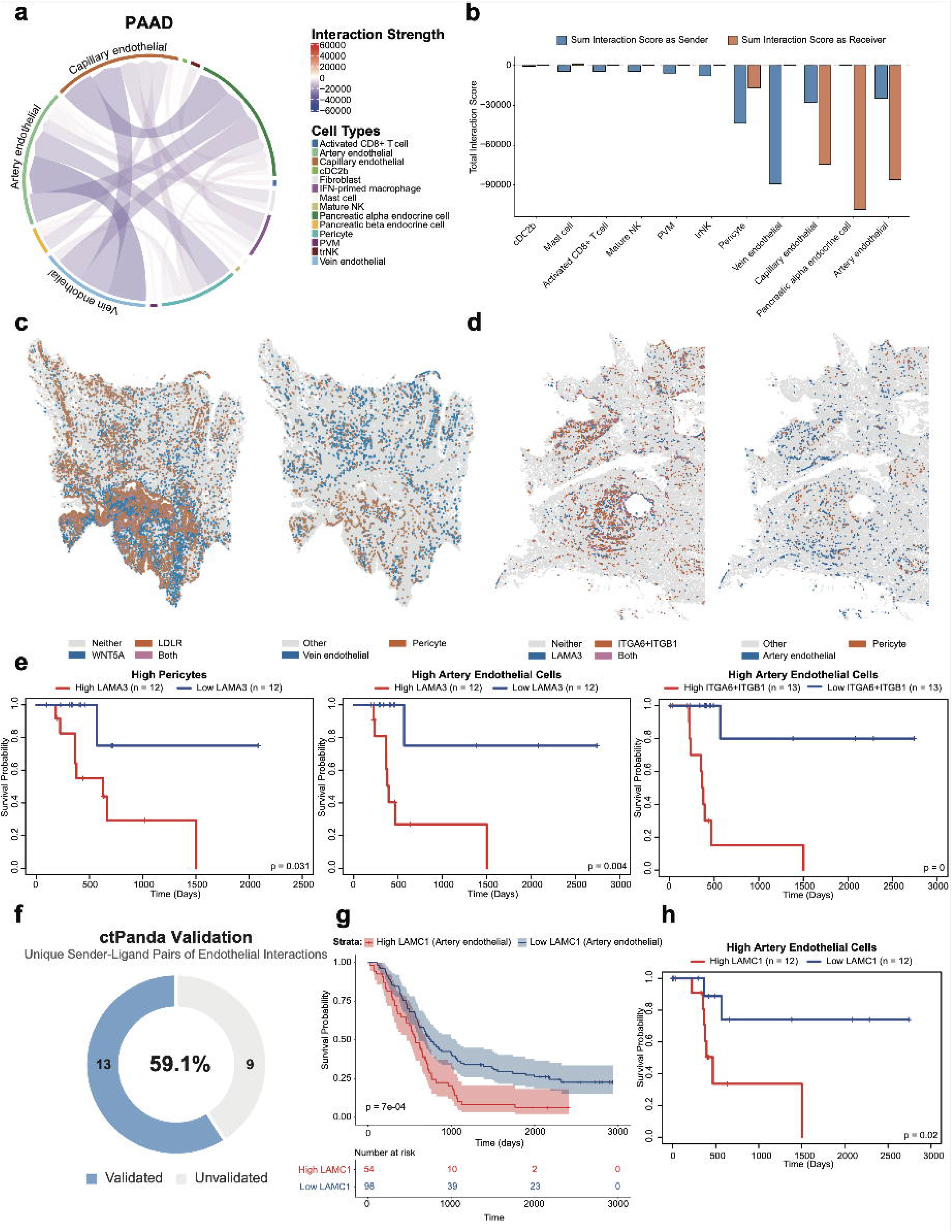
SKyNet reveals survival-associated endothelial-pericyte communication in pancreatic adenocarcinoma. **(a)** Chord diagram of SKyNet-inferred ligand-receptor interactions among cell populations in pancreatic adenocarcinoma (PAAD). Chord color indicates the survival association of each interaction (red: increased survival; purple: decreased survival) and edge color indicates cell type; endothelial populations are labeled. **(b)** Total interaction score per cell type, summed over outgoing (sender, blue) and incoming (receiver, orange) edges. **(c)** Xenium spatial transcriptomics of PAAD tissue. Left: spatial distribution of WNT5A (blue) and LDLR (orange) transcripts; regions with both transcripts are shown in pink. Right: spatial locations of vein endothelial cells (blue) and pericytes (orange). **(d)** Left: spatial distribution of LAMA3 (blue) and ITGBA6/B1 (orange) transcripts; regions with both transcripts are shown in pink. Right: spatial locations of artery endothelial cells (blue) and pericytes (orange). **(e)** Kaplan-Meier plots of overall survival of TCGA-PAAD patients stratified by expression of LAMA3 within pericyte-high patients (left), LAMA3 within artery endothelial-high patients (center), and ITGBA6/ITGB1 within artery endothelial-high patients (right). **(f)** Cross-validation of predicted endothelial sender-ligand interactions in independent PAAD validation cohort (ctPANDA). **(g-h)** Corresponding Kaplan-Meier plots of OS based on expression of LAMC1 among patient tumors enriched for artery endothelial cells by ctPANDA **(g)** and SKyNet **(h)**.

Furthermore, the spatial distribution of LAMA3 and ITGA6/ITGB1 transcripts coincided with the presence of artery endothelial sender cells and pericyte receiver cells (**Figure 4D**). Among TCGA PAAD patients whose tumors were enriched for artery endothelial cells, high LAMA3 expression was associated with significantly worse survival by Kaplan-Meier analysis, and among patients whose tumors were enriched for pericytes, high LAMA3 and ITGA6/ITGB1 expression were associated with significantly worse survival (**Figure 4E**). Because SKyNet uses bulk and single-cell RNA-seq data from different patient cohorts, we next evaluated whether SKyNet-predicted interactions could be validated in an independent PAAD cohort with patient-matched single-nucleus RNA-seq and longitudinal outcome data from 152 patients (ctPANDA)^28^. Among SKyNet-predicted interactions involving endothelial cells, 13 of 22 unique sender-ligand combinations (59.1%) showed concordant significance and poor survival outcomes in the ctPANDA cohort (**Figure 4F-H**). Therefore, SKyNet predictions in PAAD were consistent with spatial transcriptomics data and with an independent survival cohort, which suggests that SKyNet can be used to nominate novel ligand-receptor interactions implicated in cancer progression for further investigation.

### Lymphatic endothelial cell communication networks are associated with poor survival outcomes in lung adenocarcinoma

Given that SKyNet’s predicted CCC pathways in PAAD were dependent on smaller *in silico* selected cohort sizes, we also applied SKyNet to a larger TCGA lung adenocarcinoma (LUAD) patient cohort. Of the 188 interactions predicted by SKyNet (**Figure S5A, Supplemental Table 4**), interactions with lymphatic endothelial cells as the receiver were enriched, and all were associated with poor outcomes (**Figure 5A,B**). On closer examination of these lymphatic endothelial interactions, we identified putative interactions between myofibroblasts and lymphatic endothelial cells via the VEGFC-ITGB1 axis (**Figure 5C,D**). We also found putative negative interactions between lymphatic endothelial cells and neutrophils via the SPON2-ITGB1 axis (**Figure 5C,E**). SPON2 expression has previously been associated with lymph node metastasis in lung adenocarcinoma^29^, and VEGFC has been implicated in tumor cell invasion and metastasis^30^. Because interstitial macrophages and lymphatic endothelial cells were among the cell types most associated with negative outcomes (**Figure 5B**), we next focused on interactions between these two cell types, which were enriched for collagen-integrin signaling (**Figure 5F**). An independent 10x Visium spatial transcriptomics dataset of patient-matched tumor and adjacent normal tissue from 5 LUAD patients^31^ showed that co-localization of COL5A2 and the ITGA1/ITGB1 complex was significantly higher in tumor than in adjacent normal tissue from the same patient, as measured by the Jaccard similarity score (**Figure 5G, Figure S5B**). These findings are consistent with literature showing that integrin-collagen signaling can promote metastasis in lung and other cancers^32^. Together, these analyses indicate that SKyNet can reveal putative CCC pathways with reported roles in tumor metastasis in LUAD.

**Figure 5.**
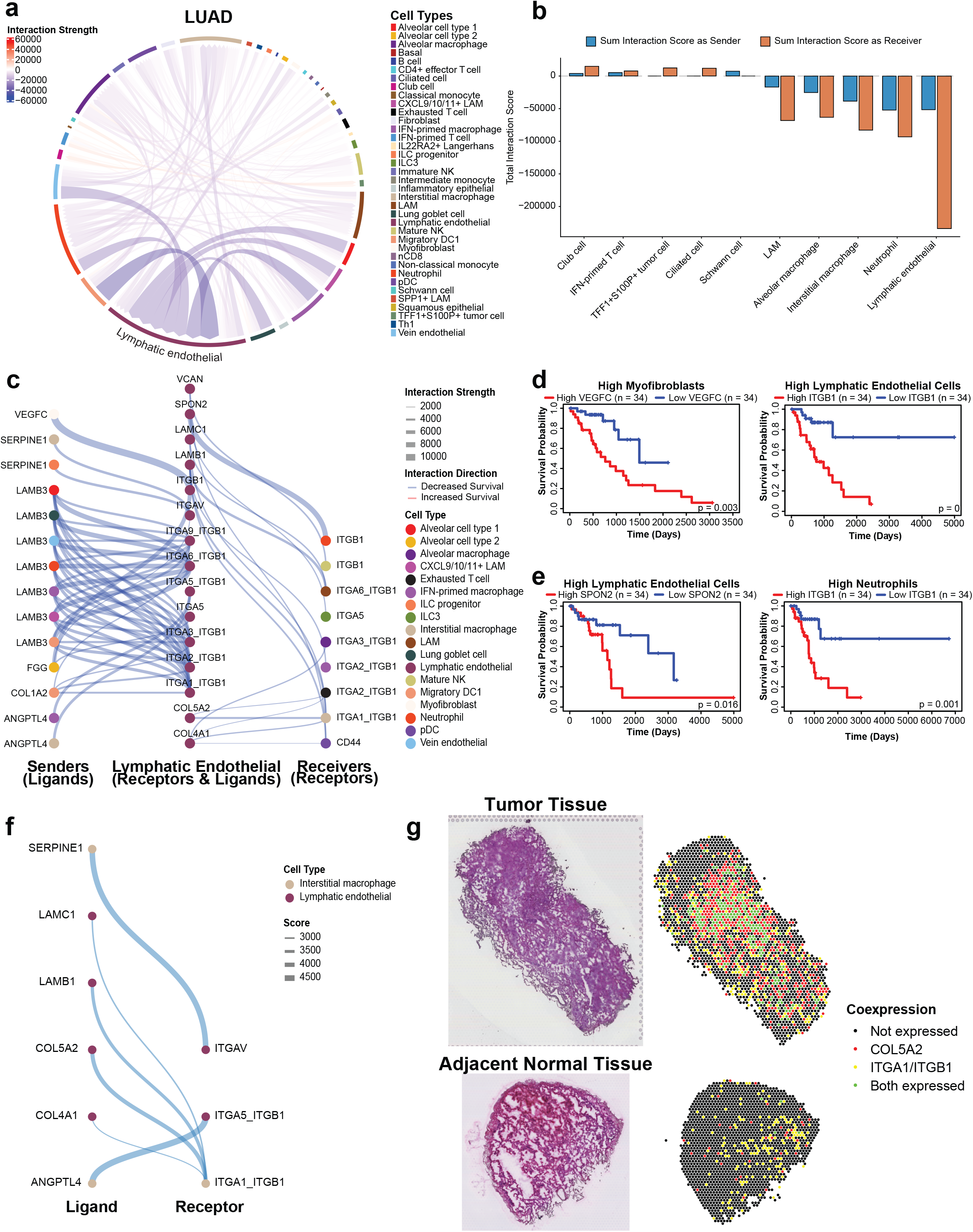
Lymphatic endothelial cell networks are associated with poor survival outcomes in lung adenocarcinoma. **(a)** Chord diagram of SKyNet-inferred ligand-receptor interactions among cell populations in lung adenocarcinoma (LUAD). Chord color indicates the survival association of each interaction (red: increased survival; purple: decreased survival) and edge color indicates cell type; the lymphatic endothelial population is labeled. **(b)** Total interaction score per cell type as sender (blue) and receiver (orange). **(c)** Lymphatic endothelial-centered communication network. Left: incoming ligands from various cell types (identified by color in legend). Center: receiving receptors on lymphatic endothelial cells and outgoing ligands on lymphatic endothelial cells. Right: Receptors on various cell types (identified by color in legend) receiving ligands from lymphatic endothelial cells. Thickness of line reflects interaction strength, and color reflects survival association. **(d-e)** Kaplan Meier survival curves of selected sender-ligand and receiver-receptor pairs VEGFC from myofibroblasts to ITGB on lymphatic endothelial cells **(d)** and SPON2 from endothelial cells to ITGB1 on neutrophils **(e)**. **(f)** SKyNet-predicted interactions between interstitial macrophages and lymphatic endothelial cells. **(g)** 10x Visium spatial transcriptomics of LUAD tissue. Spatial distribution of COL5A2 (red) and ITGA1/ITGB1 (yellow); regions with both transcripts are shown in green.

### Divergent fibroblast communication networks distinguish survival associations in subtypes of kidney cancer

To determine whether SKyNet predicted cancer CCC networks were independent of tissue-of-origin, we compared kidney renal papillary cell carcinoma (KIRP) to KIRC outputs. Indeed, the CCC landscape was distinct for each tumor (**Figure 6A,B**, **Figure 6SA,B, Supplemental Table 4**). Fibroblast communication networks showed opposite survival trends in the two cancers: fibroblast interactions were associated with prolonged survival in KIRC, whereas fibroblast interactions were mainly associated with decreased survival in KIRP (**Figure 6C,D**). In KIRC, SKyNet-predicted interactions involving DSC2/DSG2 and CDH1, which are key components of desmosomes, were associated with prolonged survival (**Figure 6C,E**). Loss of desmosome signaling has been reported to precede epithelial-to-mesenchymal transition (EMT) and to promote metastasis in certain cancers^33,34^, although its role in KIRC has not been explored. Similarly, SKyNet predicted that CDH1-PTPRM signaling was associated with prolonged survival (**Figure 6C,F**), which is consistent with previous research demonstrating that PTPRM knockdown led to larger tumors in KIRC mouse models^35^. In contrast, KIRP-predicted fibroblast interactions were dominated by integrin-collagen signaling (**Figure 6D**), which has been associated with metastasis in other cancers^36^. Several interactions between fibroblasts and immune cells were also associated with poor overall survival. For example, CCN4-ITGB1 signaling between CD8^+^ T cells and fibroblasts was associated with significantly worse survival (**Figure 6D,G**). CCN4 has been implicated in metastasis in esophageal, prostate, pancreatic, and other cancers^37^ and in immunosuppression in melanoma^38^, but it has not been studied in KIRP. SKyNet also identified LTA-TNFRSF1A signaling between exhausted T cells and fibroblasts (**Figure 6D,H**). Similarly, lymphotoxin alpha/TNF beta (LTA/TNFB) has been implicated in T cell lymphoma progression, although its role in KIRP is unknown^39^. TNFRSF1A is understudied in the context of KIRP, but it has been reported to have cancer-promoting effects in clear cell renal carcinoma cells *in vitro*, including effects on migration and invasion^40^, which suggests that a similar mechanism may operate in KIRP. Together, these findings indicate that SKyNet can identify cell type-divergent CCC associated with survival in tumors of the same organ, potentially revealing clinically relevant pathways for future investigation in human solid tumors.

**Figure 6.**
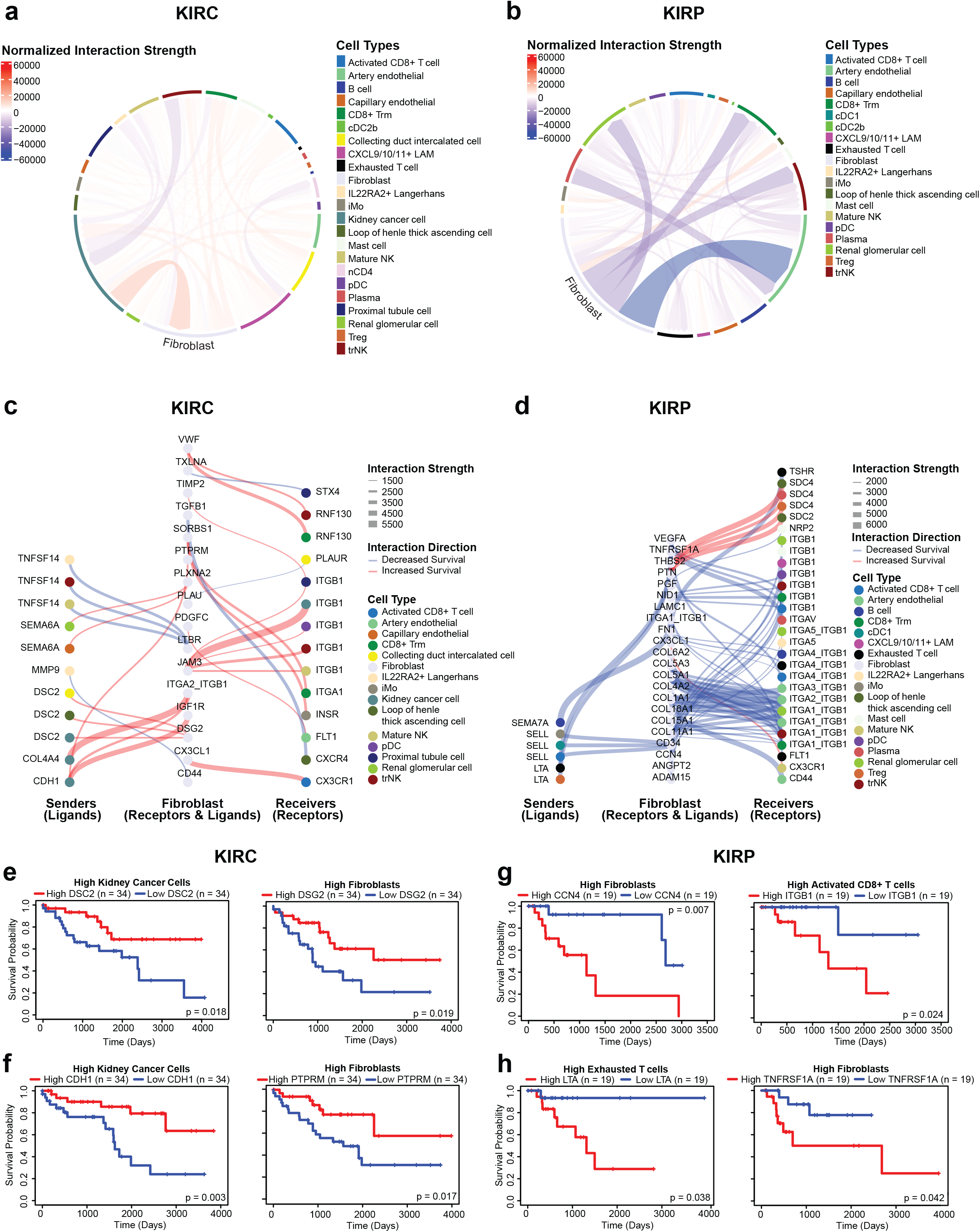
Divergent fibroblast extracellular matrix networks distinguish survival associations in KIRC and KIRP. **(a-b)** Chord diagrams of SKyNet-inferred ligand–receptor interactions in kidney renal clear cell carcinoma (KIRC) **(a)** and kidney renal papillary cell carcinoma (KIRP) **(b)**; interaction strengths normalized to enable direct cross-subtype comparison. Chord color indicates the survival association of each interaction (red: increased survival; purple: decreased survival) and edge color indicates cell type; fibroblasts are labeled in both networks. **(c-d)** Fibroblast-centered communication networks in KIRC **(c)** and KIRP **(d).** Thickness of line reflects interaction strength, and color reflects survival association**. (e-f)** Kaplan-Meier curves of the DSC2 to DSG2 interaction **(e)** and the CDH1 to PTPRM interaction **(f)** in KIRC. **(g-h)** Kaplan-Meier curves of the LTA to TNFRSF1A interaction **(g)** and the CCN4 to ITGB1 interaction **(h)** in KIRP.

## DISCUSSION

SKyNet utilizes deconvoluted bulk RNA-seq data from the TCGA patient cohort, which provides larger population sizes than are currently available from certain single-cell matched survival datasets across several solid cancers. This approach enables sufficient statistical power to predict cellular communication networks in solid cancers associated with survival following permutation testing. The primary limitation of this approach is the relatively small number of patient-matched tumor-adjacent and tumor scRNA-seq samples available for constructing high-resolution reference single-cell expression levels of ligands and receptors across solid cancers, which limited our current analysis to 11 solid tumors. However, SKyNet was able to predict communication axes cross-validated with independent spatial transcriptomics datasets, including Xenium Prime and Visium platforms, which confirmed the physical co-localization of predicted ligand-receptor pairs in independent lung and pancreatic cancer patients. Furthermore, SKyNet predicted cell types and ligands in pancreatic cancer that overlapped with an independent survival cohort^28^, which integrated single-nucleus RNA-seq from 152 patients with longitudinal outcomes. These results demonstrate that putative CCC axes can be validated across clinical cohorts even with modest scRNA-seq cohort sizes when coupled with large-scale bulk RNA-seq matched survival data.

While single-cell sequencing matched with patient outcomes has been investigated in skin, liver, and pancreatic cancers^28,41^, these studies have typically relied on integrated atlases that have unequal cellular composition across patients, complicating downstream Kaplan-Meier survival analyses without carefully balanced cohort selection. This limitation is particularly evident for rare cell populations, which are systematically underrepresented in pancreatic cancer^28^. SKyNet corrects for this limitation by selecting *in silico* patient cohorts of equal size through quartile-based cell deconvolution thresholding, ensuring that cell type composition does not limit survival comparisons when associating ligand and receptor pairs with patient survival. Furthermore, whereas ctPANDA performs pseudo-bulk analysis to correlate cell type-resolved gene expression with survival^28^, SKyNet infers ligand and receptor expression directly from single cells, enabling identification of specific intercellular communication axes rather than aggregate transcriptional associations that may be biased by single patients. Of course, observed discrepancies between these approaches may also arise from differences in cell annotation strategies, tumor sampling methods, and the inherent biological differences between single-nucleus and single-cell RNA sequencing platforms^42^, which can capture distinct cellular populations and transcriptional states. Recently, scSurvival^41^ introduced an attention-based multiple-instance learning framework that directly models patient survival from single-cell data while preserving cellular heterogeneity, demonstrating robust performance in melanoma and liver cancer cohorts. However, these approaches remain constrained by the limited cohort sizes of patient-matched scRNA-seq datasets relative to the bulk RNA-seq cohorts leveraged by SKyNet. Like these previous studies, we did not perform formal covariate analyses, such as adjusting for patient age, tumor stage, treatment history, or other clinical confounders. This was primarily due to the well-documented incompleteness of TCGA clinical metadata^14^. TCGA was designed primarily for molecular characterization, and clinical data collection was secondary, resulting in non-uniform follow-up intervals, incomplete treatment annotations, and variable reporting across tissue source sites^14^. Despite this limitation, the modular design of SKyNet permits straightforward incorporation of additional clinical covariates as more comprehensive and harmonized clinical datasets become available, and the computational framework can be readily extended to integrate treatment response data, comorbidity indices, or other patient-level variables in future analyses.

The cellular communication pathways identified by SKyNet in human solid tumors are predominantly associated with metastasis, such as stromal remodeling, immune evasion, and angiogenic signaling^43^. For example, loss of CDH1, which SKyNet predicted was associated with poor survival in KIRC, is a well-established precursor to epithelial-to-mesenchymal transition (EMT), and activation of CDH1 can reduce metastasis^44,45^. Similarly, in LUAD, SKyNet predicted that the enrichment of integrin B1^36,46^, LAMB3^47^, VEGF^48^, and SERPINE1^49^, all of which are known to play a role in local invasion in lung and other cancers, were associated with poor overall survival outcomes. However, some SKyNet-identified pathways in liver cancer may also contribute to systemic manifestations of cancer, such as cancer cachexia. For instance, CSF3, which SKyNet predicted to be associated with poor survival in HNSC (**Supplemental Table 4**), has recently been studied as a marker of cachexia in both lung and colorectal cancer^50^. Meanwhile, oncostatin M (OSM) was demonstrated to induce angiogenic effects on endothelial cells and potentially contribute to atherosclerosis^51^, which could shorten overall survival. Similarly, SERPINE1 (PAI-1) signaling has been previously implicated in venous thromboembolism, the second leading cause of death in cancer patients^52^, and in pancreatic cancer^53^. As such, SKyNet may be useful in predicting what clinical manifestations are likely to drive mortality given a patient’s identified tumor CCC network. Therefore, by defining a new class of clinically actionable cellular communication pathways that can be identified cheaply and at scale from publicly available data, SKyNet may also provide a generalizable framework for dissecting clinically relevant CCC across human disease.

## MATERIALS AND METHODS

### Data and Code Availability

All annotated Seurat objects and survival analysis code are uploaded to GitHub. The bulk RNA-seq and clinical data used in these analyses were sourced from TCGA are available in the National Cancer Institute Genomic Data Commons (GDC) Portal.^54^

### Software

All scRNA-seq analysis, survival analyses, and 10x Genomics Visium analysis were conducted using the R software^55^. 10x Xenium analysis was conducting using Python. A complete list of software, packages, and versions can be found in the key resources table.

### Single Cell Sequencing Analysis

#### scRNA-seq pan-cancer data integration and individual scRNA-seq datasets

To achieve a comprehensive representation of tissue types and cell types, we retrieved six publicly available, patient-paired scRNA-seq datasets, in addition to a non-patient-paired breast cancer dataset (**Table 1**). The numbers of patients from these datasets were selected based on a balanced total cell count. Datasets were loaded into Seurat on R and integrated using Seurat’s streamlined IntegrateLayer function and the Harmony method^56^. In addition, five other publicly available, patient-paired scRNA-seq datasets were retrieved and analyzed separately from the atlas (**Table 2**).

#### Quality control and processing of scRNA-seq data

The CreateSeuratObject function was used to import the raw data from the above sources into Seurat and conduct quality control, such as the percentage of mitochondria genes, the total number of RNA (nCount), and the number of unique RNA detected in each cell (nFeature). Cells with more than 20% of transcripts mapping to mitochondria genes and lower than 100 nCount RNA were removed. Outliers more than 2 standard deviations away from the mean of nCount and nFeature RNA were also removed. Normalization, variable features selection, data scaling, and principal component analysis were then performed. To correct for batch effects, the harmony package was used to harmonize across patient samples and conditions^6^. Harmonized cells were then clustered and visualized in Uniform Manifold Approximation and Projection (UMAP) manifold embedding.

#### Cell type annotations of scRNA-seq data

We carefully selected a list of lineage genes from the literature and existing algorithms like XCell^12^ and CellTypist^13^ for clustering and sub-clustering of all scRNA-seq datasets. Differentially expressed genes ranked by positive average log2FC from the Seurat FindMarkers function with a 25% minimum expression, were utilized along with independent interpretation for cell type identification.

To improve our annotation of organ-specific cell types, we found annotated scRNA-seq datasets of human normal tissues on the Cell X Gene atlas containing tissue-specific structural cells such as alveolar cells type 1 and 2 in the lung, cholangiocytes and sinusoidal endothelial cells in the liver, and granulosa cells and theca cells in the ovary^57^. We also referenced a comprehensive breast cancer atlas for breast-specific cell types^58^. Lineage genes for these cell types were retrieved using the FindMarkers function and ranked by positive log2FC.

#### Distinct module scores of cell types

For each cell type, we defined transcriptional modules as the top 20 genes ranked by positive average log2FC in the integrated scRNA-seq object (**Supplemental Table 2**). As with our lineage genes, we further curated these modules by manually adding canonical genes from existing atlases and algorithms.

We then validated that each cell type could be transcriptionally distinguished through their module score. We used the AddModuleScore function in Seurat to calculate the module score of each cell, comparing module expression to background expression. Kruskal-Wallis testing evaluated whether the module scores revealed significant differences between cell types. Subsequently, median module scores were normalized and scaled to generate z scores and to distinguish if each cell type is represented the best with its corresponding gene module.

#### Lineage markers for Individual scRNA-seq datasets

From the integrated atlas, we defined a list of module scores reflective of cell type transcriptional modules across cancers, which we called common lineage markers. For each individual scRNA-seq object, we also identified cancer-specific cell types and generated transcriptional modules for them by taking the top 20 genes ranked by positive average log2FC. The common lineage markers and cancer-specific cell type markers were utilized in downstream survival analyses.

#### Area under precision-recall curve (AUPRC)

To evaluate the specificity of our transcriptional modules, we generated precision-recall curves for each cell type across 12 annotated scRNA-seq datasets using the ROCR package^59^. We calculated the mean area under the curve of each precision-recall curve across cell types for each cancer. We conducted the same analysis with the transcriptional modules from XCell^12^ (FANTOM and BLUEPRINT) and CellTypist^13^ (Immune Encyclopedia Version 2) to facilitate comparison with existing these algorithms.

#### Single-cell percent and scaled average normalized expression

For each scRNA-seq dataset, percentage of expressing cells and scaled average expression per cell type of all genes were determined from raw count matrices.

For multi-protein receptor complexes, a cell was defined as expressing the complex only if all constituent genes were detected (expression > 0). For positive cells, complex expression was defined as the mean expression of its component genes; cells lacking any required gene were assigned zero expression. Percent expression was calculated as the proportion of cells within a cell type expressing all constituent genes. Average expression was quantified by taking the z-score of the scaled average expression across all cells within that type.

#### Survival Analysis

25 different human solid cancers from TCGA were used for lineage-based survival analysis: BLCA, BRCA, CESC, COAD, ESCA, GBM, HNSC, KIRC, KIRP, LGG, LIHC, LUAD, LUSC, OV, PAAD, PCPG, PRAD, READ, SARC, SKCM, STAD, TGCT, THCA, THYM, and UCEC. Of these, 11 were used to infer significant ligand-receptor interactions (CESC, COAD, GBM, HNSC, KIRC, KIRP, LIHC, LUAD, LUSC, OV, and PAAD). These cancers were chosen based on scRNA-seq dataset availability, which allowed us to extract lineage markers for tissue-specific cell types. Overall survival was used in all survival analyses, which was defined as days until the last follow up or days until death. Survival analyses were performed using Survival for cox proportional hazards models and Survminer for Kaplan Meier analysis.

#### Processing of Bulk RNA-seq & Clinical Outcome Data

Bulk RNA-seq and clinical outcome data were extracted from TCGA’s GDC Portal via the TCGABiolinks package^60^. Bulk RNA-seq data were processed using the DESeq2 package^61^. Genes with fewer than 10 counts across all patients were excluded, and data were normalized using variance stabilizing transformation. Patients without primary tumor samples were excluded, as were those who did not have available outcome data. A table of sample sizes for each TCGA dataset used is available in **Supplemental Table 3**.

#### Single-Lineage Survival Analysis

To perform single lineage-based survival analysis, patients were ranked based on their expression of the lineage marker gene modules derived from previous scRNA-seq analysis. For a given cell type, each patient was ranked for each gene in the lineage marker set, where the top rank (1) was assigned to the patient with the highest expression. Once each patient had been ranked for each gene, each patient was given a mean rank, which was calculated as that patient’s average rank across all genes in the module. Subsequently, the patients in the top quartile for mean rank were selected as the “*High*” group, while the patients in the lower quartile were selected as the “*Low*” group. After integrating with clinical outcome data for each patient, a univariate Cox proportional hazards model was fitted using the survival package^62^. The forest plot visualization was created using the forestplot package^63^. P-values were calculated by performing permutation testing analysis of the adjusted hazard ratios using base R. Heatmap visualization was created using the pheatmap package^64^.

#### Lineage-Ligand Survival Analysis

Lineage-ligand analysis was performed to determine the stratification power of specific ligands and receptors. For a given cell and ligand (or receptor) patients were first grouped into “*High*” and “*Low*” groups based on their expression of the cell’s lineage markers. Then, the upper quartile of patients was re-ranked based on their abundance of the ligand. Within this sub cohort, the upper quartile of patients based on ligand expression was labeled the “*High/High*” group, indicating high abundance of both cell and ligand, while the lower quartile of patients based on ligand expression was labeled the *“High/Low”* group, indicating high abundance of cell but low abundance of ligand. Then, Kaplan-Meier (KM) analysis was performed between the *“High/High”* group and *“High/Low”* group. The survival score of for sender *i* and ligand *l* was calculated as follows:

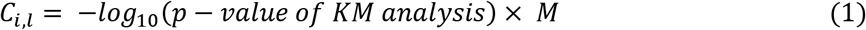

where *M* is the difference in restricted mean survival time and is calculated with:

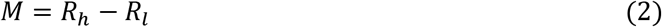

where *R_h_* and *R_h_*are the restricted mean survival times of the *“High/High”* and *“High/Low”* groups, respectively.

Representative survival curves were generated using the survminer package in R and p-values calculated based on the KM analysis^65^.

#### Identification of Significant Interactions

To identify clinically significant interactions, we ran lineage-ligand analysis for every represented cell-ligand and cell-receptor pair in 11 human solid cancers. Ligand-receptor interactions were drawn from scDiffCom, a curated database of ligand-receptor pairs in humans^20^. Using scRNA-seq data, we filtered for interactions in which one cell (the sender) expressed the ligand and the other cell (the receiver) expressed the receptor for each ligand-receptor pair; a cell type was said to express a gene if greater than 5% of cells of that cell type expressed that gene based on scRNA-seq data. We further filtered these interactions by keeping only those interactions for which receiver-ligand, receiver-receptor, and sender-ligand survival analyses showed significant (*p ≤ 0.05*) differences that were in the same direction (e.g. the survival difference (Equation 2) was > 0 for all pairs or < 0 for all pairs).

#### Filtering of Biomarkers

Biomarkers were defined as ligands and receptors whose KM analysis showed a significant difference for at least two-thirds of cell types. Given their lack of specificity, ligand-receptor interactions involving biomarkers were removed. The complete filtering procedure is visualized in Supplemental Figure 2.

#### Permutation Testing

To ensure robustness of our final predicted ligand-receptor interactions, we conducted permutation testing of all predicted cell/ligand and cell/receptor pairings (referred to as the “pair” henceforth). This was done as follows: first, the baseline survival score (Equation 1, 2) from KM analysis was calculated for a given pair; then, for each of 1000 iterations, the patients’ overall survival values were randomly reshuffled and the new survival score was calculated given this randomly shuffled data; finally, the permutation p-value for the pair was defined as the number of iterations out of 1000 in which the survival score from the randomly permuted data was greater than or equal to the baseline survival score. Pairings with a permutation *p-value ≥ 0.05* were eliminated from SKyNet’s final predictions.

#### Visualization and Sender Prioritization

For each receiver-receptor-ligand combination, there were several potential senders based on scRNA-seq data. To prioritize the top sender for a particular combination for visualization purposes, we calculated a sender score for each sender *i* and ligand *l* as follows:

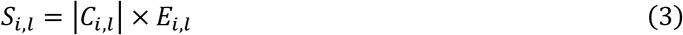

where *C_i,l_* is the survival score (Equation 1, 2) of the lineage-ligand analysis for sender *i* and ligand *l*. *E_i,l_* is the normalized expression of ligand *l* in sender *i* based on scRNA-seq data. Subsequently, the sender was chosen using:

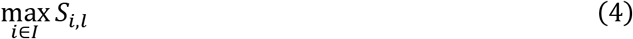

where *S_i,l_* was the set of sender scores for all possible senders *I* of ligand *l*. A flow chart of these steps can be found in.

To compare different ligand-receptor interactions across cell types, we computed an interaction score for a receiver i, ligand l, and receptor r as:

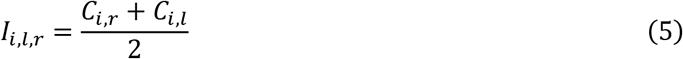

In instances where the same receiver cell *i* received the same ligand *l* through a set of multiple receptors *R*, we summed the interaction scores for each receptor to create a total interaction score for that receiver-ligand pairing as:

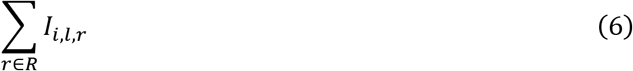

Subsequently, in these instances, we colored the dot for that pairing according to the receptor that had the greatest interaction score. These results were visualized using ggplot2 package^66^.

To assess the strength of different pairs of interacting partners, an interaction strength score for receiver *i* and sender *j* was calculated as:

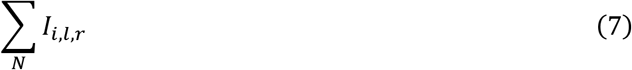

where *N* represented all the predicted interactions between receiver *i* and sender *j* where *i* expressed receptor *r* and *j* expressed ligand *l*. To visualize these interactions with a chord diagram, we used circlize package^67^.

#### CellChat and NicheNet

To compare SKyNet results to pre-existing algorithms, we ran CellChat and NicheNet on the same 11 scRNA-seq datasets using the default settings for each.

To run CellChat, we used the CellChat package on R^2^. Human signaling interactions (CellChatDB.human) were calculated using the triMean statistical method (computeCommunProb) and filtered to interactions present in at least 10 cells per group. Communication probabilities at both the pathway and aggregate levels were computed. Tumor and adjacent tissue were processed separately and then merged using the mergeCellChat function. Fold changes were calculated using a pseudo count of 1×10^−5^. Interactions were filtered to remove autocrine signaling (same receiver and sender) and homotypic (same ligand and receptor) interactions. Candidates were filtered with a minimum base communication probability of 10% and a 100-fold change. For each receptor-target pair, the primary sender displaying the highest absolute fold-change was selected.

To run NicheNet, we used the nichenetr package on R^3^. NicheNet ligand-receptor databases were prefiltered to exclude predicted but unvalidated protein-protein interactions from the ppi_predictions_go and ppi_prediction databases. Expression was filtered at a minimum threshold of 5%. For each cell type, differential gene expression analysis between tumor and adjacent tissue-derived cells were calculated using Wilcoxon rank-sum tests using the FindMarkers function, which requires at least 3 cells per condition. Differentially expressed genes (adjusted p < 0.05 and | log2FC | > 0.25) were used to predict ligand activity within each condition. Predicted ligand interactions were scored and standardized into Z-scores per receiver cell type and condition. Interactions exceeding a Z-score threshold of 2.5 were retained.

For all three networks (NicheNet, CellChat, and SKyNet), identified interactions were classified based on the cell type identity (immune or stromal) of the sender and receiver cell types. Interactions were annotated using Biomni based on the biological function of the ligand-receptor interaction (**Supplemental Table 5**)^21^. Venn diagrams were generated using the ggvenn package^68^.

#### ctPanda

To evaluate robustness of our predicted pancreatic adenocarcinoma (PAAD) interactions in an independent cohort, we generated Kaplan-Meier survival curves using ctPanda, a publicly available resource^2828^. Patients were stratified as previously described^28^. Survival curves and log-rank test p-values were generated for all unique sender-ligand pairs identified by SKyNet.

Spatial Transcriptomics

#### 10x Xenium

We analyzed publicly available 10x Genomics Xenium spatial transcriptomics data from six patients with pancreatic adenocarcinoma (PAAD)^22^. The 10x Xenium data was loaded into Scanpy in Python^69^. After filtering out cells with fewer than 10 transcripts or 5 unique genes, we conducted standard quality control, entailing normalization, variable gene selection, and principal component analysis. To reduce batch effects, the samples were then harmonized by using the harmonypy package^6^. Following batch correction, a k-nearest neighbor graph was constructed using the harmonized principal components (n = 30 components, 15 neighbors). Cells were then clustered by the Leiden algorithm and visualized in the UMAP embedding. As with scRNA-seq, cell type annotations were determined using curated lineage markers from literature.

To evaluate the degree of clustering within ligand-receptor pairs predicted by SKyNet, we performed Moran’s I analysis using the SpatialDM package^23^. Spatial weight matrices were generated using a scale parameter l=1.2 and a spatial cutoff of 0.2. Global and local spatial co-expression statistics, alongside significant spot identification, were calculated using permutation testing (n=1,000 iterations, *p < 0.05*).

#### 10x Visium

We analyzed publicly available 10x Genomics Visium spatial transcriptomics data of paired tumor and adjacent tissue slices from five lung adenocarcinoma (LUAD) patients^31^. 10x Visium data was processed in R using Seurat, following the same computational pipeline and software versions as described for the scRNA-seq analysis.

Expression scores for ligand and receptor gene modules were calculated for each spot using the AddModuleScore function on Seurat. Spots with a module score above zero were classified as positive for that gene module. Each spot was categorized into four distinct groups based on module expression: both, only Module 1, only Module 2, or none. Categorized spots were overlaid onto corresponding H&E histological images using the SpatialPlot function.

To quantify the spatial colocalization of ligand-receptor interactions, we calculated Jaccard similarity (J) between ligand and receptor expression for each spot:

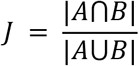

where A and B represent the set of spots positive for Module 1 and Module 2, respectively. Differences in Jaccard similarity between paired adjacent and tumor tissue across patients were evaluated using paired two-tailed t-tests.

## ACKNOWLEDGEMENTS

T.E.O. is supported by the National Institutes of Health (NIH) (R01AI174519, R01AI186079) and the UCLA Immunology Advisory Committee. Our results are based on data from The Cancer Genome Atlas, which was supported by the National Cancer Institute (NCI) and the National Human Genome Research Institute (NHGRI). We are grateful to De Zuani et al. (2024) and Wainberg et al. (2026) for releasing their datasets for public use.

## AUTHOR CONTRIBUTIONS

T.E.O. conceived and designed the project, supervised experiments, and wrote the manuscript. D.L., V.B., and J.L. analyzed data, generated figures, and wrote the manuscript. G.R.L. and J.H.J. generated figures and supervised experiments. E.C. and K.H. analyzed data.

---

**Supplemental Figure 1. Merged atlas harmonization and JadeCell module score evaluation on scRNA-seq datasets. (a-b)** Uniform manifold approximation and projection (UMAP) of all cells colored by cancer type **(a)** or patient **(b)**. **(c)** Enrichment of JadeCell module scores across all 78 cell types identified in the pan-cancer scRNA-seq atlas. **(d)** Mean area under the precision recall curve (AUPRC) across cell types for JadeCell (dark green), CellTypist (light green), and xCell (blue) gene modules in 12 cancers using t-testing. Data represents mean ± SEM. *p < 0.05, **p < 0.01, ***p < 0.001; ****p < 0.0001; ns, not significant.

**Supplemental Figure 2. Pre-processing and integration of data from The Cancer Genome Atlas.** Pre-processing and integration pipeline of clinical and RNA-sequencing data from The Cancer Genome Atlas (TCGA). Figure created with BioRender.

**Supplemental Figure 3. Ligand-receptor database comparison and interaction family classification of NicheNet- and CellChat-predicted interactions. (a)** Overlap of ligand-receptor pairs in the NicheNet, CellChat, and SKyNet databases. **(b-c)** Functional classification of **(b)** NicheNet and **(c)** CellChat interactions across cancers by Biomni (Huang et al., 2026). Each L–R pair was assigned to one of 16 functional classes; point size denotes interaction score, and color denotes the direction of the survival association (blue, decreased survival; orange, increased survival).

**Supplemental Figure 4. SKyNet-predicted interactions and 10x Xenium spatial transcriptomic validation of endothelial interactions in pancreatic adenocarcinoma. (a)** Dot plot of SKyNet predicted ligand-receptor interactions in pancreatic adenocarcinoma (PAAD). Each point represents a predicted ligand-receptor pairing; dot size is proportional to the interaction score and significance of survival; square points denote interactions associated with increased survival, and round points denote interactions with decreased survival. Sender score is the survival score of the Kaplan-Meier analysis of the sender-ligand pairing, and the absolute value is plotted. **(b)** Cross-validation of predicted endothelial ligand-receptor interactions using spatial co-localization in Xenium spatial transcriptomics data of patient-matched normal and cancerous pancreatic tissue.

**Supplemental Figure 5. SKyNet predicted interactions and 10x Visium spatial transcriptomic validation of collagen-integrin interactions in lung adenocarcinoma. (a)** Dot plot of SKyNet predicted ligand-receptor interactions in lung adenocarcinoma (LUAD). Each point represents a predicted ligand-receptor pairing; dot size is proportional to the interaction score and significance of survival; square points denote interactions associated with increased survival, and round points denote interactions with decreased survival. Sender score is the survival score of the Kaplan-Meier analysis of the sender-ligand pairing, and the absolute value is plotted. **(b)** Jaccard similarity index of COL5A2 and ITGA1/ITGB1 spatial co-localization in five patient-matched tumor and adjacent spatial transcriptomics samples of LUAD.

**Supplemental Figure 6. SKyNet predicted interactions in KIRP and KIRC. (a-b)** Dot plot of SKyNet predicted ligand-receptor interactions in kidney renal clear cell carcinoma (KIRC) **(a)** and kidney renal papillary cell carcinoma (KIRP) **(b)**. Each dot represents a predicted ligand-receptor pairing; dot size is proportional to the interaction score and significance of survival; square dots denote interactions associated with increased survival, and round dots denote interactions with decreased survival. Sender score is the survival score of the Kaplan-Meier analysis of the sender-ligand pairing, and the absolute value is plotted.

