## Supplementary figures and images for "Survival-associated cellular communication networks in human solid cancers"

### Supplemental Figure 1

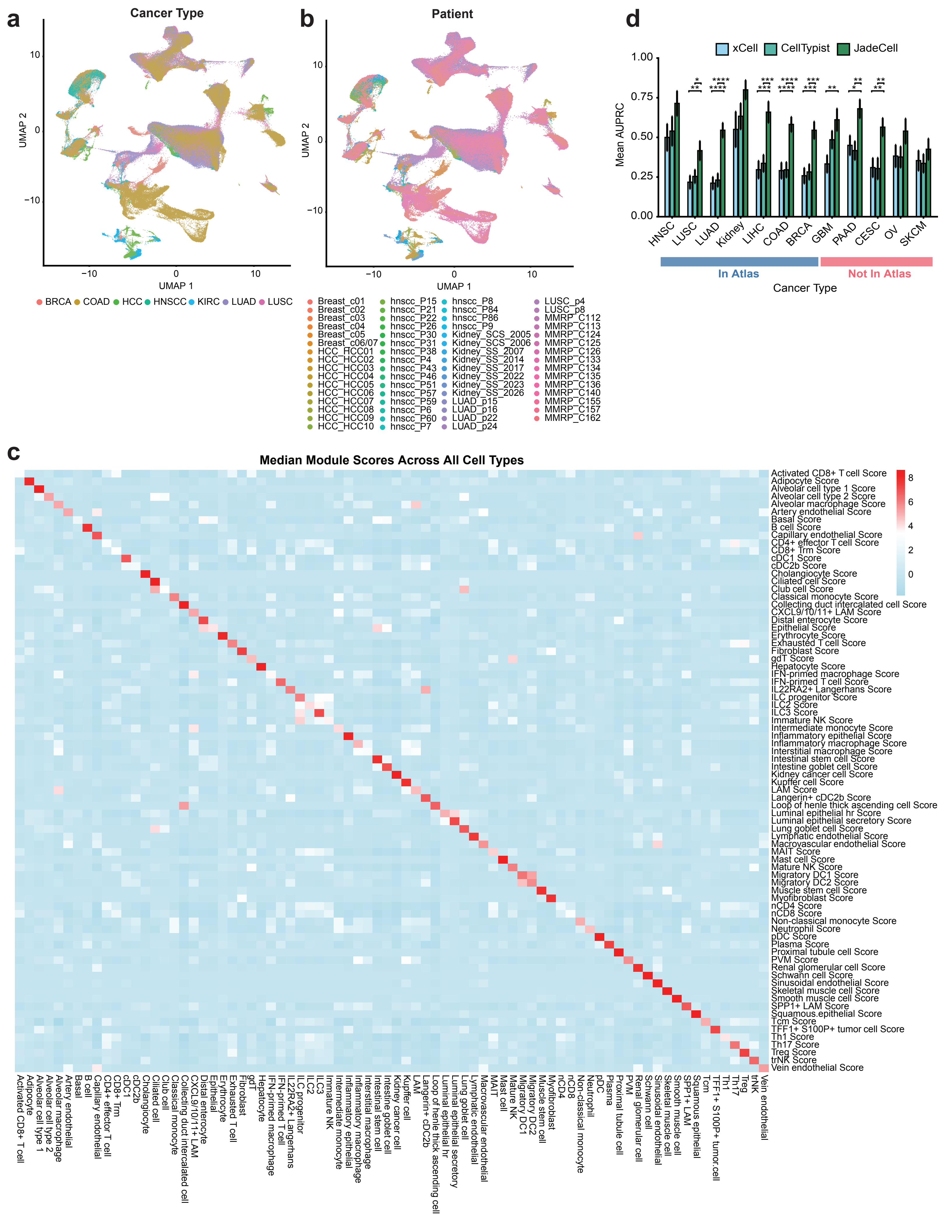

### Supplemental Figure 2

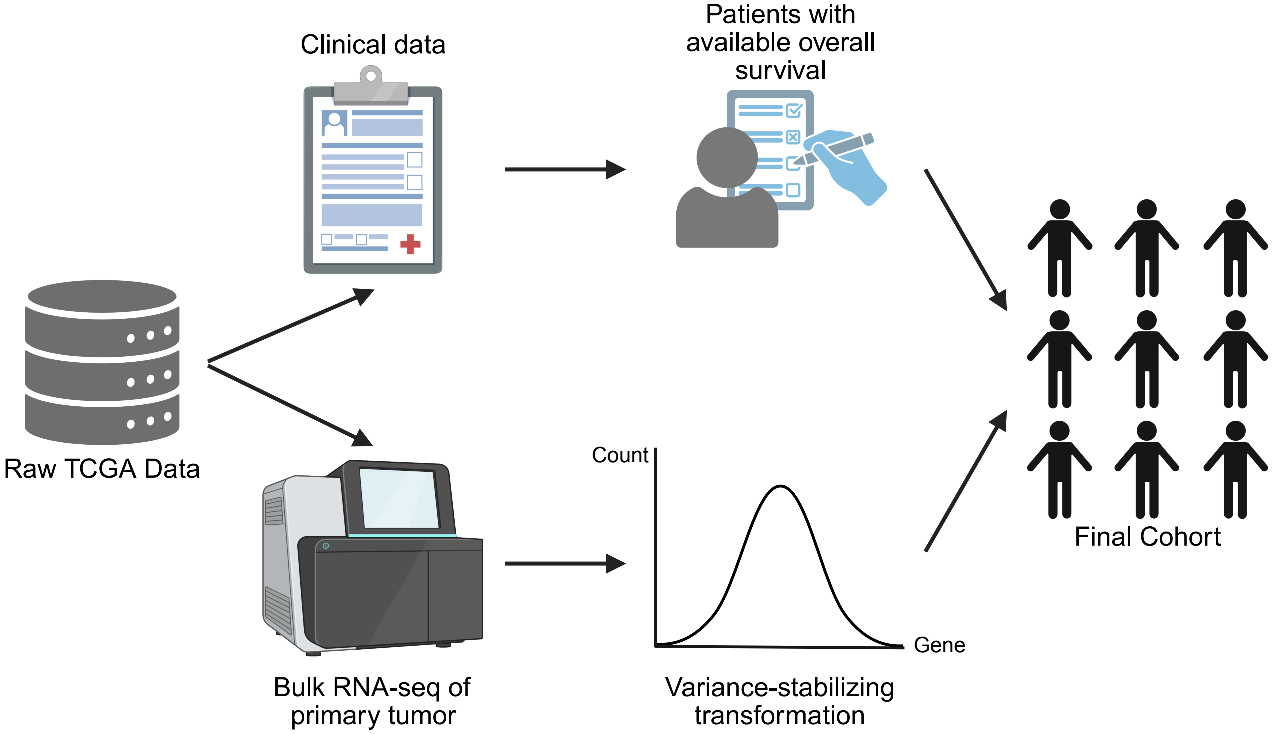

### Supplemental Figure 3

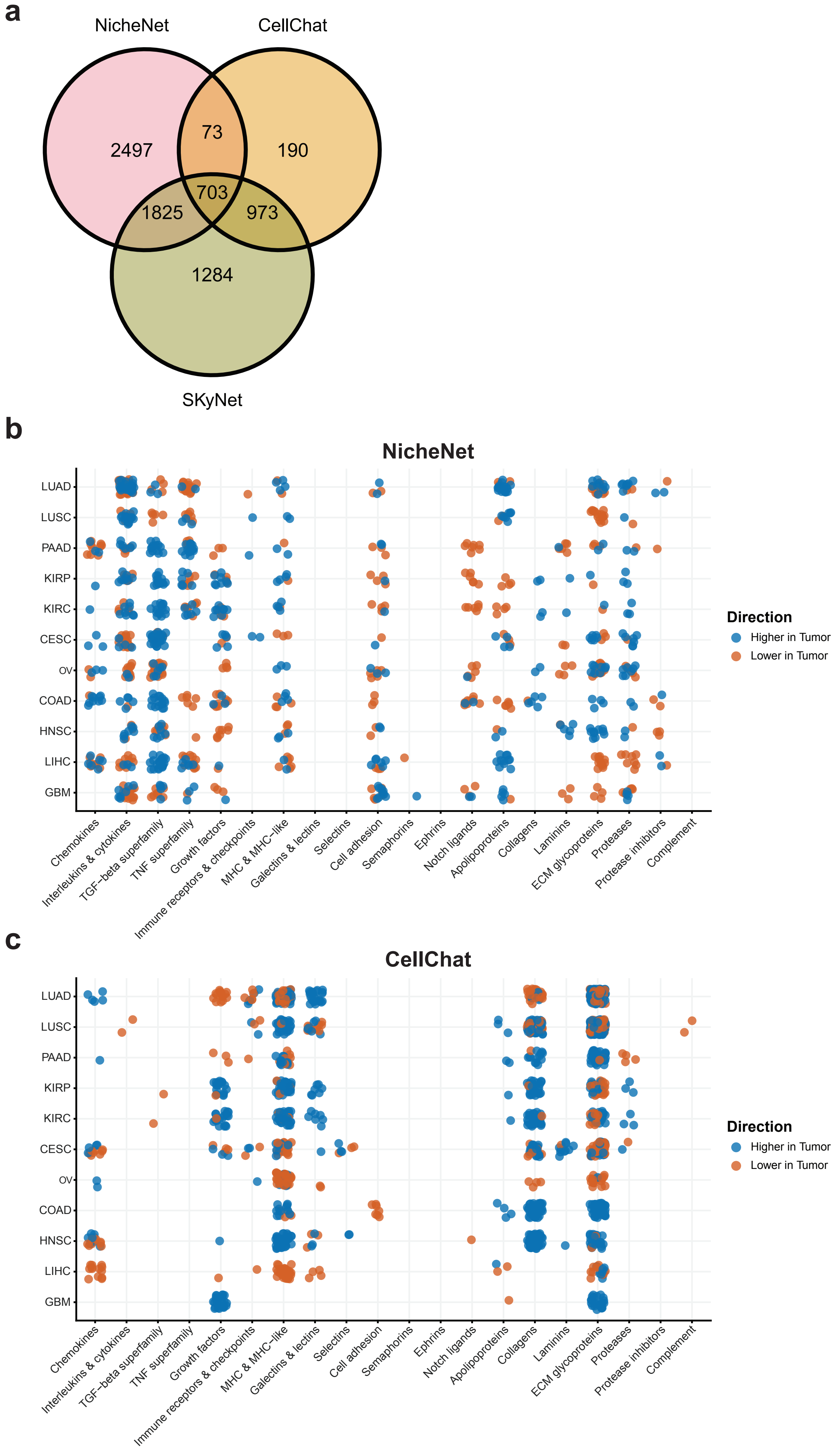

### Supplemental Figure 4

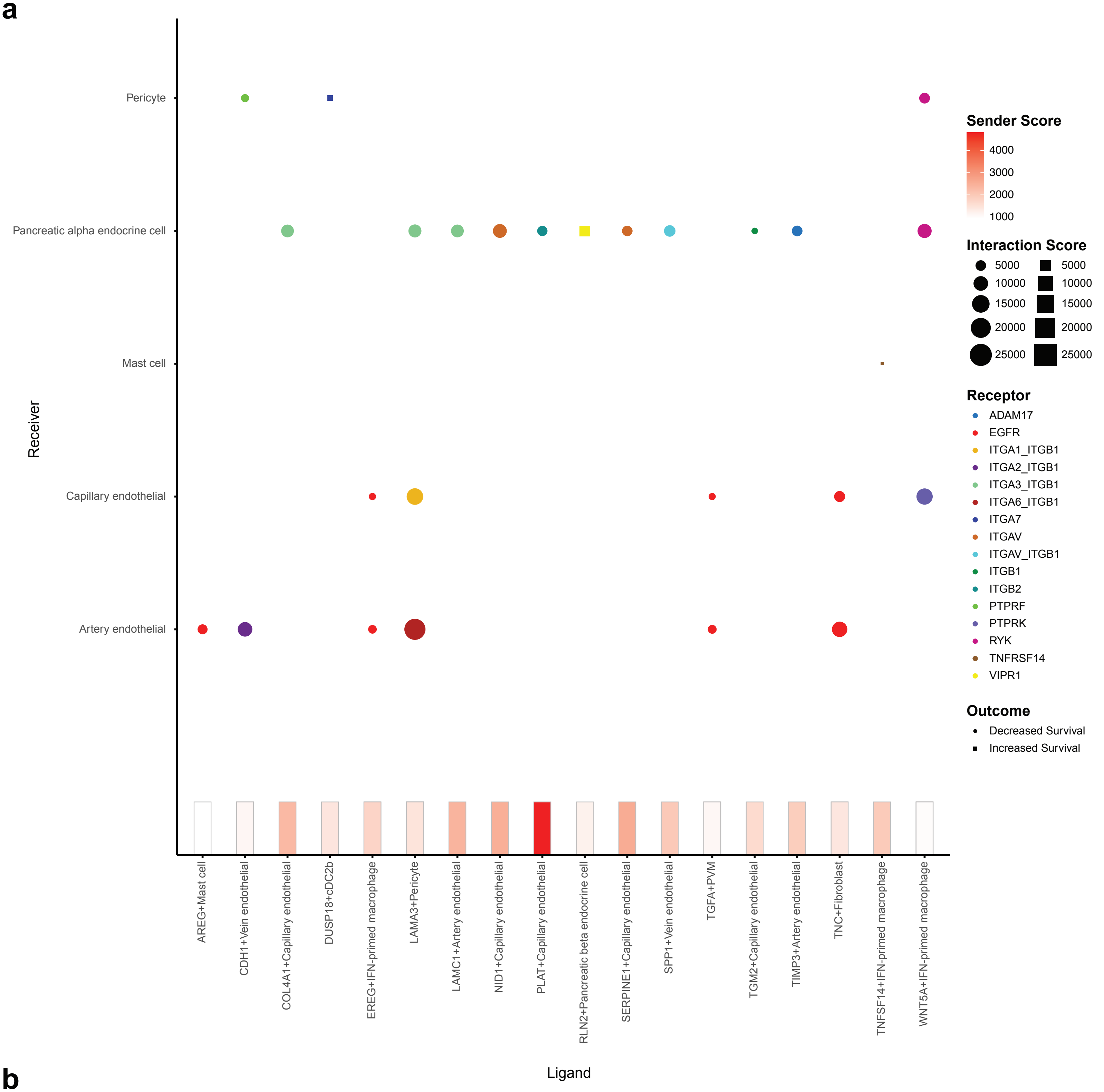

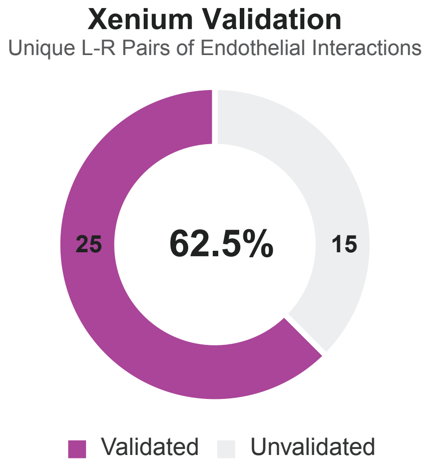

### Supplemental Figure 5

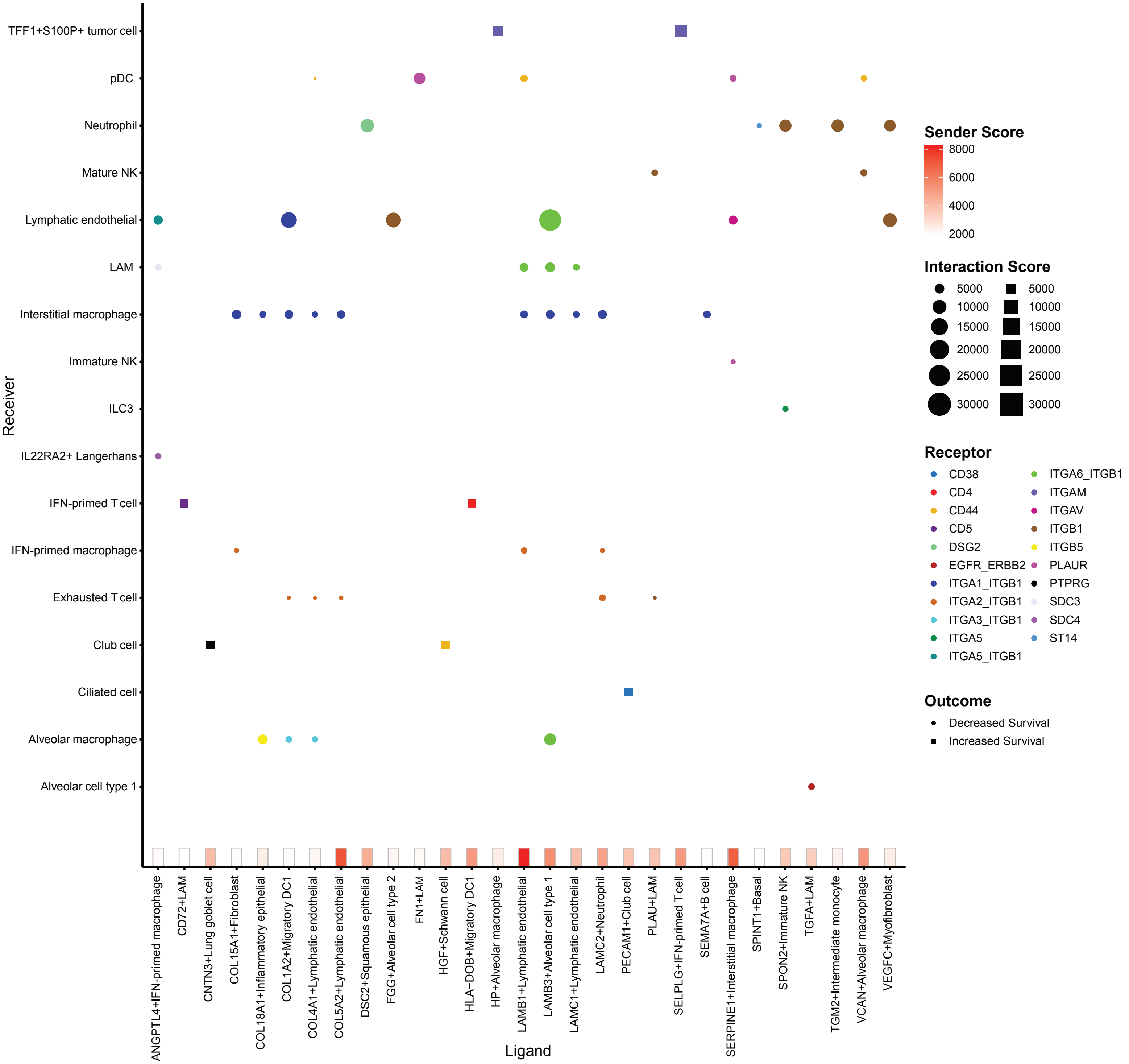

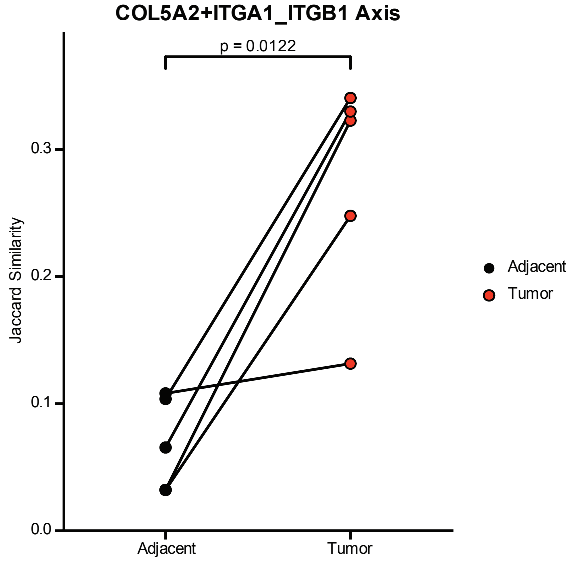

### Supplemental Figure 6

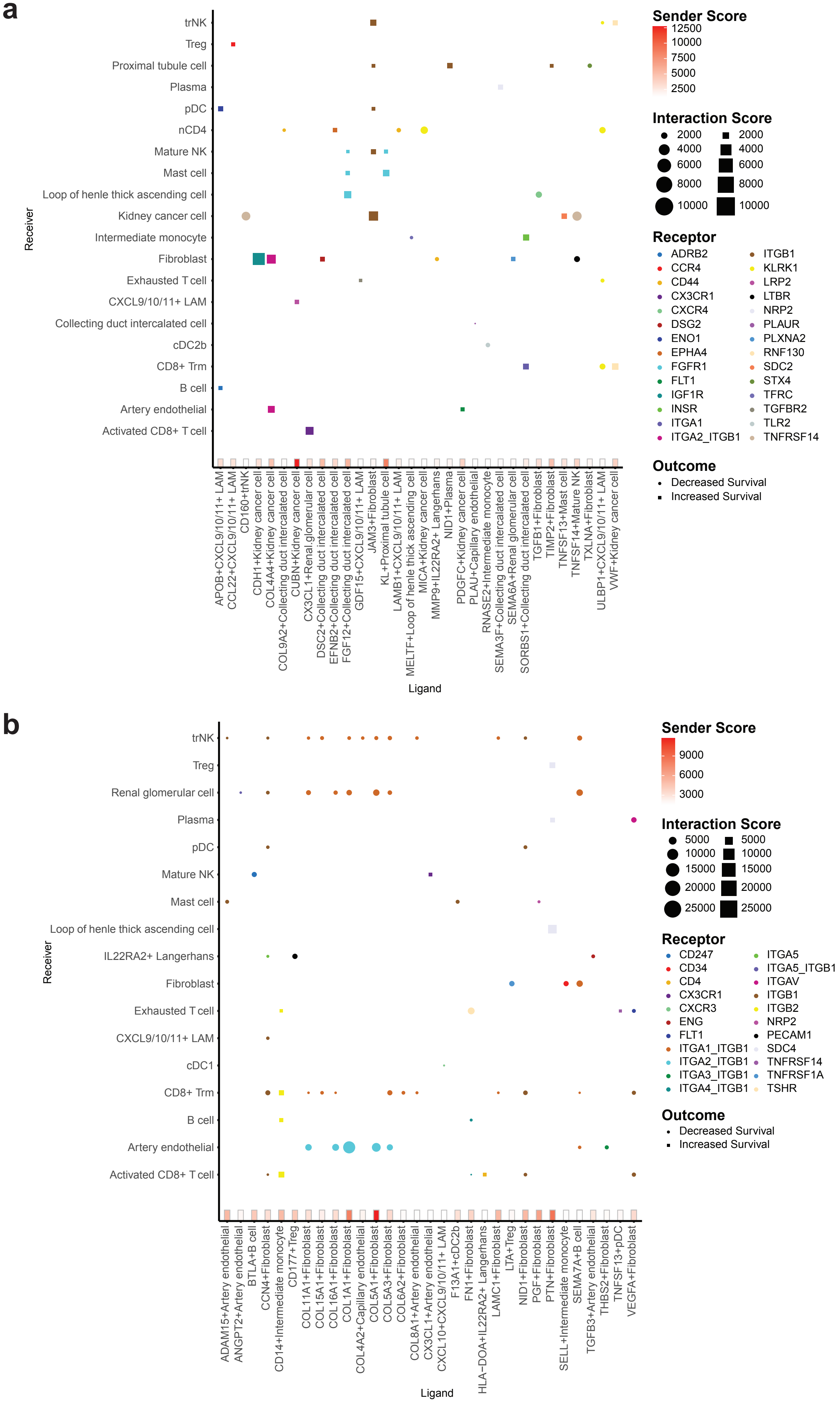
